# Human exploitation of shellfish in the Atacama desert coast and environmental variability: a trans-Holocene perspective

**DOI:** 10.64898/2026.02.02.703213

**Authors:** Bernardo R. Broitman, Laura Olguin, Javiera Guardia, Mauricio H. Orostica, Adrien Chevallier, Luna Vasquez, Carola Flores

## Abstract

The Humboldt upwelling ecosystem has been intensively harvested by people since the early Holocene. Understanding past and present human choices under climatic variability in these productive environments may hold key insights for its future sustainability by unraveling different adaptive pathways. To this end, we studied shellfish exploitation and climate patterns in the Taltal region of the Atacama desert coast (25^◦^S) from the early Holocene until today using a compilation of archaeological, and modern benthic fisheries data together with direct ecological surveys. In addition we obtained satellite sea surface temperature (SST) and published *δ*^18^O SST for the study region. The archaeological record and the modern rocky shore assemblage were dominated by herbivorous gastropods –*Fissurella* spp., *Enoplochiton* spp., *Tegula* spp.–and the carnivorous whelk *Concholepas concholepas*. Functional composition from the early Holocene to the present was remarkably stable. Using SST as a latent variable, we examined changes in functional composition across the Holocene and in a 16-year series of artisanal fisheries landings using bayesian ordination. The analysis identified functional groups characteristic of kelp ecosystems in association with cooler SST conditions during the Holocene and the present. Changes in functional composition during warm and cold periods of the Holocene broadly mirrored effects of interannual SST variability in the modern fisheries. The archaeological record suggests two cross-Holocene transitions social-ecological transitions. The generalized shoreline harvesting strategy that prevailed during the cold early Holocene shifted to a specialized maritime economy towards the warmer mid-Holocene. The maritime technological and cultural adaptions remained, but were part of more diversified lifestyles in the cooler and more variable late Holocene. The latter emerged at the same time as the modern El Niño climate pattern. Our insights from the direct analysis of human choices and SST variability highlight the role of flexibility and agency under a changing environment. The broad range of human decisions in the past, inform current regulatory frameworks for benthic artisanal fisheries. Marine resources and the livelihoods that depend on them are integrated into coupled coastal socioecological systems; their future sustainability hinges on fostering the different dimension of their adaptive capacity.

## 1 Introduction

### 1.1 Human exploitation of coastal resources and environmental variability

Human populations crossed the Bering strait during the late Pleistocene and arrived to South America during the Pleistocene-Holocene transition (12500-10000 cal yr BP)^91,74^. Terrestrial habitats were long regarded as the main pathway for the early peopling of the Americas; it is now understood that coastal environments played key role along the initial stages of the process^45,44^. An extreme case is the Atacama region. The driest desert in the world, it extends between the Pacific ocean and the Andean highlands, from 18^◦^ to 27^◦^S^43^. Few permanent rivers drain into sea^107^ and the narrow and mostly rocky littoral platform is flanked by a tall and steep coastal range (Figure 1). The coastal ocean, however, is the northern part the Humboldt current eastern boundary upwelling ecosystem^105^. Following the scarcity or remoteness of terrestrial resources, the earliest inhabitants of the Atacama coast were heavily reliant on the coastal environment for subsistence^102,70,101^. Coastal upwelling along the Atacama coast is driven by semipermanent equatorward wind along the eastern Pacific margin –and the rotation of the earth that steer surface waters offshore–to bring deeper, colder, and nutrient rich waters to the surface^105^. The recently upwelled waters fuel the striking primary and secondary production that characterizes the Humboldt region^6,35^. Over interannual scales, the El Niño-Southern Oscillation (ENSO) cycle modulates the efficiency of upwelling by changing the flux of nutrients to the sunlit surface as warm (cold) water is advected poleward (equatorward), thus deepening (shoaling) the thermocline^11,109,46^. As such, ENSO-driven changes in marine productivity have extensive impacts on the region’s coastal social-ecological systems in the present^11,34,35,23^. Similarly, archaeological evidence has shown that the livelihoods of human societies inhabiting the Atacama coast in the past where impacted by climatic-scale modes of environmental variability, including ENSO-like changes to the SST regime ^101,99^. The impact of past climate changes on coastal societies in the Atacama has been well documented through archaeological shell middens, which chronicle their livelihoods^81,82,69,50^. The identity and relative abundance of different remains in archaeological shell middens is a legacy of past human choices, technology, and the environmental context where shellfish were collected^93,94,121^. On the other hand, independent geochemical proxies from archaeological material or geological deposits can help reconstruct past environmental conditions on multiple spatial and temporal scales^110,27,50^. When combined, they can provide a compelling background to discern to what extent the composition and abundance of archaeo-faunal assemblages were driven by changes in environmental conditions, socio-cultural decisions, or a mixture of both^27,67,49,100,69^. Today, the impacts of climate-driven variability are being compounded by deep anthropogenic impacts to marine food webs^88,77^. Learning the lessons from past climate impacts on coastal social-ecological systems can provide important insights for the future^47^.

**Figure 1:**
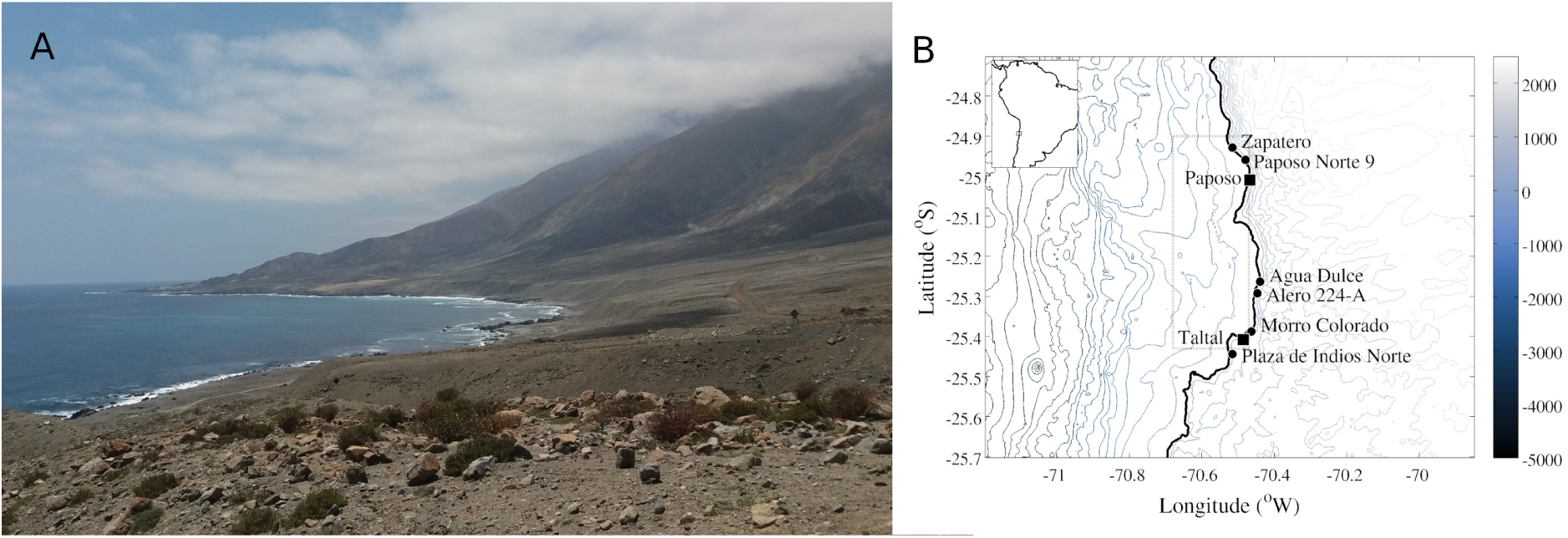
**A**. Image of the Taltal coastline from Alero 224-A looking equatorward. Notice the aridity of the landscape and the steep topography of the coastal range. Road sign at the right of center for scale. **B**. Topographic map of the Taltal area showing the steep bathymetry of the coastal zone –depths in excess of 5000 are located 30-40 kilometers offshore. The location of the six archaeological shell midden sites are indicated with black circles, the harbors of Taltal and Paposo are indicated with black squares, and the dotted polygon offshore corresponds to the area from where we extracted and averaged the MODIS satellite SST time series for statistical analyses. The small inset on the upper left indicates the study area location on the coast of western South America.Topographic data from^56^

### 1.2 Archaeological and Paleoceanographical patterns across the Holocene

Our work focuses on the Taltal area (25^◦^S, Figure 1), where a chronocultural sequence has been proposed to organize the long human record of the region based on radiocarbon dates and stratigraphical characteristics of the archaeological evidence^96^. During the early Holocene (Archaic I, 12000-10000 cal yr BP) cold sea surface temperature (SST) nearshore are interpreted as intense upwelling conditions^53,86,30,50^. Archaeological sites for the period are characterized by few shell midden deposits beneath small rock shelters, with fire pits and remains of domestic occupations and located within 200 meters of the modern coastline. Cultural remains indicate the use of terrestrial, coastal and marine resources; lithic artifacts were made of raw materials obtained from the adjacent inland ^97^. Around 10000 cal yr BP, marine transgressions flooded 1-2 km of coastline submerging archaeological sites near the shore and part of the early Holocene settlement pattern^96^.

Hence, a period of 1500 years lacks archaeological remains. The next open-air residential site is dated to the early-mid Holocene (Archaic II, 8500 cal yr BP) and the evidence from the initial occupation suggests short-term but recurrent visits, with debris from domestic and workshop activities^7,96^. At the beginning of the Middle Holocene (Archaic II-II, 8500-6900 cal yr BP) inshore SST was as warm as present at 25^◦^S^52,50^. Warmer SST inshore and offshore may be linked to a weak seaward temperature gradient and curtailed upwelling, which is supported by different paleoceanographic proxies^73,98,50^. After around 7500 cal yr BP, the stratigraphic sequence of the latter open-air residential site shows a gradual change toward a more dense and long-term occupation, which marks the beginning of a specialized maritime economy^96,51,92^. Between 7500 and 5500 cal yr BP (Archaic III), there is an increase in the size and number of open-air shell midden sites with high abundance and diversity of artifacts and ecofacts. During the latter part of the mid-Holocene, between 6000 and 4300 cal yr BP, isotopic analyses in archaeological shells from the Taltal area show a decrease in SST with the coolest moment of the late Holocene around 4300 cal yr BP associated with depleted *δ*^13^C values, suggesting an increase in upwelling activity^50^. Importantly, the onset of the modern ENSO pattern has been proposed for the second half of the mid-Holocene^99,110,119,29^. Around 5500 yr BP (Archaic IV), changes in the settlement system proposed as evidence of territoriality^32,10,79,89,96^.

Finally, paleoceanographic evidence indicates that during the Late Holocene (c.a. 4000 cal yr BP onward), increased ENSO activity was associated to higher and more variable SST and weaker coastal upwelling in southern Peru and northern Chile^86,29,30,50^. After 4500 cal yr BP (Archaic V) the number of sites along the Taltal coast decreased; the accompanying cultural changes suggest regional transformations related to more diversified subsistence economy, together with changes in fishing technology and cultural norms^32,96,4^. Onward from 3500 cal BP (Archaic VI), there is a noticeable increase in the number of residential and funerary sites in the area^97^. Despite evidence of cultural and material exchange with far-inland communities, the use of maritime and coastal resources remained constant. Together, the changes and adaptation in the social-ecological system across the Holocene for the Taltal region suggest a strong feedback between SST conditions, resource availability and cultural patterns.

### 1.3 Coastal ecology and small-scale fisheries in the present

Along the cold and temperate Pacific coast of the Americas, different laminarial algal species form massive subtidal and intertidal stands –kelp forests– that support a characteristic assemblage exploited in the past and in the present^45,92,81,90,26^. The diverse and productive kelp forest communities are dominated by a few trophic-functional groups, chiefly sea urchins, herbivorous and carnivorous gastropods, and benthic carnivorous fish and seastars^41,54,19^. Changes in SST related to environmental variability can influence the extent and compositions of kelp forests, and the associated fauna, from local to regional scales^41,72,20,75,80,14,76^. Over broad latitudinal gradients, patterns of SST variability are defined by large-scale circulation regimes, which along eastern boundary ecosystems determine biogeographic structure ^17,118,48,68^. For example, the semipermanent upwelling regime along the Atacama desert coast transitions progressively to a more seasonal and the a late-season pattern poleward, which is reflected in the functional composition of the assemblage targeted by small-scale benthic fishers^36^. Similarly, upwelling regime has been shown to modulate the performance of management areas for benthic resources along central-northern Chile^8^. Small-scale fishers’ landings and effort are linked to both socio-economic and environmental drivers^40,120^. However, small-scale fisheries in Chile –artisanal fishers– are regulated through top-down management strategies^57,36^; they are allowed to operate only in the administrative region were they are registered, which curtails the adaptive capacity of the social-ecological system, particularly under a changing climate^120,38^. Earlier syntheses of paleoceanographic and archaeological information from the Atacama coast have concluded that ocean environmental transitions have coincided with shifts in the cultural trajectories of its first coastal societies^101,100,71^. Our analysis attempts to bring together the extensive archaeological record of the Taltal region together with palaeotemperature estimates in order to understand how environmental regime and human choices shifted subsistence patterns along the region ^96,92^. At the same time, we draw on nearly two decades of publicly-available artisanal landing records and satellite SST to the drivers of the modern social-ecological system to shed light on potential adaptive strategies to cope with future climate change ^38^.

## 2 Methods

### 2.1 Archaeological Molluscan assemblage

Shellfish archaeological data comes from six study sites located along the coast of Taltal (Figure 1 and Table 1). The extensive archaeomalacological dataset presented here was derived over a decade of fieldwork and analyses^82,83,84,85,111,62^. Cultural materials indicated that the main use of our study cites was as chiefly residential, with differential intensity of occupation^96^. Excavations followed cultural and natural layers, and all shellfish material was collected from 50 by 50 cm column samples ranging between 50 and 100 cm of depth.

**Table 1:** Geographic position of the six study sites together with the abbreviation used in the text and the type of archaeological deposit. Shell middens correspond to open-air shell mounds, while rock shelters are sites with small shell deposits beneath a rocky roof.

| Site name | Code | Lat°S | Lon°W | Midden type |
| --- | --- | --- | --- | --- |
| Zapatero | ZP | -24.928 | -70.516 | Shell midden |
| Paposo Norte 9 | PN9 | -24.958 | -70.480 | Rock shelter |
| Agua Dulce | AD | -25.262 | -70.439 | Shell midden |
| Alero 224-A | 224A | -25.292 | -70.447 | Rock shelter |
| Morro Colorado | MC | -25.387 | -70.463 | Shell midden |
| Plaza de Indios Norte | PIN | -25.443 | -70.516 | Shell midden |

All recovered specimens were quantified and identified to the lowest possible taxon – chiefly at the species level – using species identity catalogs and reference collections for the study area. To quantify shellfish abundance, we used Minimum Number of Individuals (MNI) for each taxon^31,16,60,93^. Before data analysis we standardized MNI values by excavated volume using the area and depth sampled for each site and chronocultural period (Supplementary Table 1, MNI / m^3^)^60,93^. For qualitative and quantitative analyses, we grouped the archaeomalacological record from the six sites using the early, middle and late Holocene or following the six chronocultural stages proposed by Salazar et al. ^96^.

### 2.2 Modern benthic shellfish assemblage

The modern shorelines of the Taltal area are predominantly rocky with limited sandy areas. Therefore, we focused our examination on macroinvertebrates to the intertidal zone of the rocky shores directly adjacent to thearchaeological sites (localities) north of the city of Taltal, between the Morro Colorado and Zapatero sites (Figure 1, Table 1). Between 2016 and 2022 and over seven visits to the area, we completed a total of 14 days of direct ecological surveys. During morning and/or afternoon low tides, we conducted 42 different 10-15 m long swath transect surveys covering, a total of 1252.4 m². We identified all mobile macroinvertebrates (i.e. larger that 1 cm) in the mid- and low-intertidal zone within a two-meter wide band centered on the transect line where we counted all individuals to estimate species density as the number of individuals per species per m². The intertidal zone around the archaeological sites of PIN and MC was not surveyed; both sites are heavily impacted by commercial and recreational shellfish gathering due to their proximity to populated areas and the intertidal zone is almost devoid of macroinvertebrates (Author’s personal observation). The rest of the sites were located on remote and uninhabited coastlines (Figure 1A), yet we always observed people gleaning for shellfish in the vicinity of the survey areas. All mollusks from ecological surveys were identified to the species level in the field except some limpets of the genus *Scurria* – *S. variabilis, S. araucana, S. ceciliana* –recent molecular genetic advances have shown that they are indistinguishable by shell morphology alone^87^. In addition to field surveys of the rocky intertidal zone, we obtained data for artisanal landings of benthic organisms compiled by scientific observers of the National Institute of Fisheries (Instituto de Fomento Pesquero – IFOP); data collection methods are fully described in^36^. Briefly, observers recorded multispecific fisheries using a standardized protocol that recorded landings from benthic capture from areas that were not subject to a management regime. We used the recorded landings of the two small fishing ports within our study area (Paposo and Taltal) that were fished along the coastline shown in Figure 1^36^. Multi-species landings used in the analysis were available as monthly landings in each port (tons) and covered a 16-year period between 2004 and 2019. As we wanted to examine temporal changes in modern fisheries with temporal variation in archaeological assemblages, we used for statistical analysis the sum of all yearly landings per fisheries group over both landing ports as a compromise (see 2.4).

### 2.3 Oceanographic and Palaeoceanographic context

To assess on the influence of environmental variability on benthic species, we focused on SST patterns^35,37,108,49,72,76,80^. For the period covered by the modern fisheries data, we used 8-day Level 3 (L3) nighttime composite data (2003-2019) SST from the Moderate Resolution Imaging Spectroradiometer (MODIS)) with a 4×4 km spatial resolution obtained from the NASA Ocean Color repository (http://oceancolor.gsfc.nasa.gov/). The L3 images were spatially and temporally averaged over the offshore polygon shown in Figure 1, corresponding to the region used in^52^ to cross-validate *in situ* and satellite SST measurements with the *δ*^18^O SST of carbonate samples from shells of modern keyhole limpets (*Fissurella maxima*). Following this validation, we utilized the trans-Holocene *δ*^18^O SST data obtained from archaeological *F. maxima* samples from^50^ as a proxy of paleoenvironmental conditions conditions in the study area.

### 2.4 Data Analysis

To explore human gathering patterns throughout the Holocene we assessed changes in abundance of individual species across the Archaic periods. The MNI of all archaeological invertebrates were grouped into functional groups based on a combination of taxonomic class and trophic level similar to the benthic fisheries groups for coastal Chile described in Chevallier et al. ^36^. We used the same functional group approach to examine the temporal variation in landing data, which were pooled over time as described in section 2.2. In addition, and in order to approximate kelp abundance in the archaeological record, we used the MNI of *S. scurra*, an exclusive epibiont of kelp, as a proxy of prehistoric kelp harvesting^4,2^.

To examine the relationship between oceanographic conditions and the abundance of different functional groups in the archaeological record and the landings data, we used SST statistics for each chronocultural period from the *δ*^18^O SST data in Flores and Broitman ^50^ and MODIS SST for modern data (Figure 1). We then characterized the variation in the composition of benthic fisheries and identified indicator taxa using a model-based approach of unconstrained ordination. We fitted a latent variable model using Bayesian Markov chain Monte Carlo estimation^66,116^ using the R package *boral* ^65^, where species’ responses were assumed to follow a Tweedie distribution with a log-link function connecting the linear predictor and the mean of the distribution. For both archaeological and fisheries data, we fitted the latent variable models using all possible combinations of SST statistics, namely the mean and the low and high temperature ranges using the 5, 50, and 95*^th^* percentiles of all the SST data available for each period or year, respectively (Figure 2). In the case of modern fisheries data, we used SST values of the previous year to reflect the lagged relationship between environmental variability and catch in harvested stocks in the Humboldt current ecosystem^6^.

**Figure 2:**
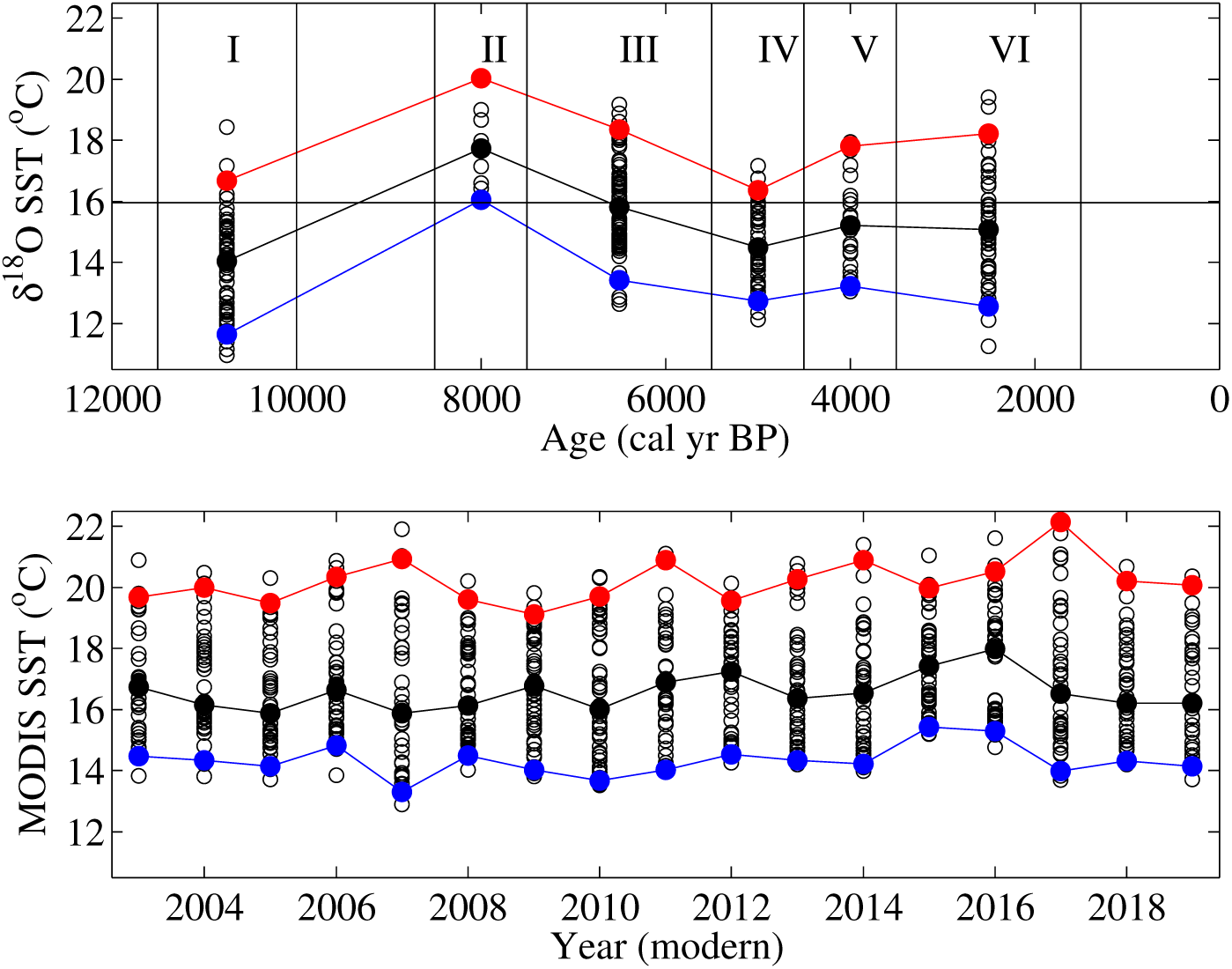
Time series of SST utilized for the ordination analyses of fisheries for the Holocene and the present. Upper panel: 5*^th^* (blue), 50*^th^* (black), and 95*^th^* (red) percentiles of the *δ*^18^O SST values observed for each of the chronoarcheological periods defined in^96^. The SST values for each period correspond to all records for individual shells presented in^50^. The solid line across the plot, around 16*^o^*C corresponds to the long-term Holocene *δ*^18^O SST mean. Lower panel: Time series of L3 MODIS SST interpolated to monthly values over the offshore section indicated by a dotted polygon in Figure 1, the same area utilized for cross-validation in^52^. Using the same criteria used for archaeological shells, for each year of the MODIS SST we selected a lower, central, and upper percentiles of a normal distribution (5, 50 and 95%) among all the monthly SST values for each year. Note that the statistical analysis for modern fisheries used SST values for the previous year.

## 3 Results

### 3.1 Archaeological, ecological and fisheries assemblages

In the six archaeological sites, we identified a total of 57 taxa, while in the adjacent rocky intertidal zone we found a total of 32 macroinvertebrate species. A total of 18 species were shared between archaeological and modern intertidal assemblages. The full list of species and abundances is provided in Supplementary Table 2. The species exclusively in the archaeological samples were mostly poorly preserved specimens identified to a lower taxonomic resolution (e.g. Order Lottiidae) or for groups that are restricted to either sandy or subtidal benthic environments. The limited taxonomic resolution in archaeological specimens may hinder direct comparisons with modern samples, as some species may in fact be shared between both assemblages but remain unrecognizable in the archaeological record due to fragmentation or erosion. Species exclusively present in ecological surveys were small crustaceans, sea stars, or species identified only at the genus level in the archaeological assemblage (e.g. Supplementary Table 2). Among the ten more common species in the archaeological record, ranked by their abundance during the early Holocene, all were relatively abundant in the modern intertidal zone assemblage. Following this ranking, the species shared between shell middens and current ecological surveys were all herbivorous gastropods (*Enplochiton* spp., *Scurria* spp. *Fisurella* spp. and *Tegula* spp.), except for the carnivorous gastropod *Concholepas concholepas* (Table 2). Other edible invertebrates that were common in ecological surveys and observed across all archaeological periods were the edible sea urchin *Loxechinus albus* and different species of Chiton (*C. granosus, C. latus, Tonicia* spp.). Species that are not edible owing to their small body size were also abundant and present in all archaeological periods were *Echinolittorina peruviana, Marinula pepita* and *Prisogaster niger*. The more abundant species only present in modern rocky shore assemblages were *Tetrapygus niger, Heliaster helianthus* and *Chiton barnesii*. Carnivorous seastars (i.e. *H. helianthus, Meyenaster gelatinosus* and *Stichaster striatus*) were also common on the rocky shore, but it was not possible to identify their presence in archaeological shell midden samples.

**Table 2:**
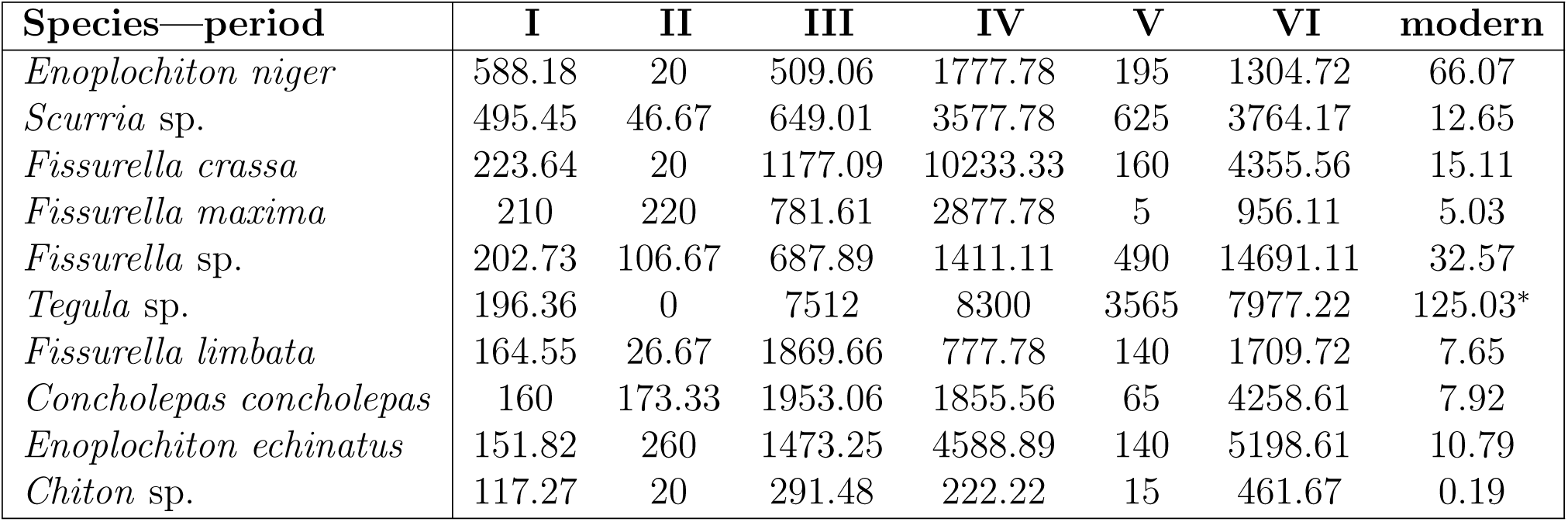
Abundances for the ten more common species in the archaeological record for each chronocultural period (MNI) and in ecological field surveys (individuals per m^2^) of the intertidal zone of rocky shores adjacent to the shell midden sites. Species are sorted using their shared rank abundance during the early Holocene. As all individuals in the modern sample were identified to species level, we present the density of *Tegula atra* in the modern sample, which is indicated by a ^∗^.

| Species—period | I | II | III | IV | V | VI | modern |
| --- | --- | --- | --- | --- | --- | --- | --- |
| *Enoplochiton niger* | 588.18 | 20 | 509.06 | 1777.78 | 195 | 1304.72 | 66.07 |
| *Scurria* sp. | 495.45 | 46.67 | 649.01 | 3577.78 | 625 | 3764.17 | 12.65 |
| *Fissurella crassa* | 223.64 | 20 | 1177.09 | 10233.33 | 160 | 4355.56 | 15.11 |
| *Fissurella maxima* | 210 | 220 | 781.61 | 2877.78 | 5 | 956.11 | 5.03 |
| *Fissurella* sp. | 202.73 | 106.67 | 687.89 | 1411.11 | 490 | 14691.11 | 32.57 |
| *Tegula* sp. | 196.36 | 0 | 7512 | 8300 | 3565 | 7977.22 | 125.03* |
| *Fissurella limbata* | 164.55 | 26.67 | 1869.66 | 777.78 | 140 | 1709.72 | 7.65 |
| *Concholepas concholepas* | 160 | 173.33 | 1953.06 | 1855.56 | 65 | 4258.61 | 7.92 |
| *Enoplochiton echinatus* | 151.82 | 260 | 1473.25 | 4588.89 | 140 | 5198.61 | 10.79 |
| *Chiton* sp. | 117.27 | 20 | 291.48 | 222.22 | 15 | 461.67 | 0.19 |

The MNI of all carnivorous gastropods such as *Acanthina monodon, C. concholepas, Crassilabrum crassilabrum* and *Xanthochorus crassidiformis* were one order of magnitude less abundant than herbivorous gastropods (Table 3). During field surveys we did not record the presence of conspicuous filter feeder species of large body size that can form dense aggregations in the shallow subtidal or low intertidal zone and were present in the archaeological record: the edible barnacle *Austromegabalanus psicattus* and mytilid mussels (i.e. *Mytilus* or *Perumytilus*). As our surveys did not target subtidal soft-bottom habitats, it was not possible to evaluate the presence of clams, bottom feeding gastropods, scallops or slipper-limpets, all common in the archaeological assemblage. In contrast, groups of species, such as decapods (crabs), and the edible sea urchin *L. albus* were common both in archaeological and modern samples (Table 3).

**Table 3:** Summed MNI for every fisheries functional group across each archaic period (I, II, III, IV, V, VI) and the field densities per m. ^2^ **of the main functional groups detected in the rocky intertidal zone adjacent to the excavated shell middens. Modern kelps densities are for individual holdfasts**^112^.

| Group—period | I | II | III | IV | V | VI | modern |
| --- | --- | --- | --- | --- | --- | --- | --- |
| Herbivorous Gastropods | 2250.91 | 980 | 9654.01 | 27911.11 | 2010 | 36359.17 | 228.48 |
| Carnivorous Gastropods | 187.27 | 180 | 2031.48 | 3455.56 | 65 | 5677.22 | 7.92 |
| Sea Urchins | 37.27 | 33.33 | 443.13 | 1111.11 | 265 | 3416.67 | 134.14 |
| Kelp | 356.36 | 2900 | 31624.68 | 10977.78 | 4195 | 19731.94 | 125.03 |
| Barnacles | 19.09 | 6.67 | 64.9 | 100 | 10 | 249.17 | 0 |
| Decapods | 10 | 20 | 58.26 | 511.11 | 15 | 384.17 | 0 |
| Bottom feeding Gastropods | 20 | 66.67 | 298.72 | 0 | 95 | 1199.17 | 0 |
| Slipper-limpets | 9.09 | 0 | 33.52 | 0 | 0 | 123.61 | 0 |
| Mussels | 0 | 60 | 223.4 | 22.22 | 0 | 335.83 | 0 |
| Clams | 9.09 | 20 | 157.17 | 0 | 0 | 62.5 | 0 |
| Scallops | 0 | 20 | 84.19 | 0 | 0 | 23.61 | 0 |
| Carnivorous seastars | 0 | 0 | 0 | 0 | 0 | 0 | 26.09 |

The IFOP data for modern fisheries landings were dominated by kelp, both subtidal and intertidal species (mainly *Lessonia* spp.); the tonnage was two orders of magnitude higher than the rest of the landings (Figure 3). Switching in between the second and third place and oscillating in the amount of total landings where sea urchins (*L. albus*) and herbivorous gastropods (*Fissurella* spp.) (Figure 3). The fourth most abundant group were carnivorous gastropods, chiefly *C. concholepas*. Except for a few years (i.e. 2006-2007), bivalve landings were exceedingly small – less than 1 ton annually – with a similarly curtailed abundance pattern for filter-feeding slipper limpets (Calypterids). It is worth noting that barnacle landings are not presented or used in the ordination analyses: only two months in the entire series reported landings of this edible species. Other groups that are present in the archaeological dataset, such as bottom feeding gastropods, scallops and decapods are absent from the groups recorded in the landings data.

**Figure 3:**
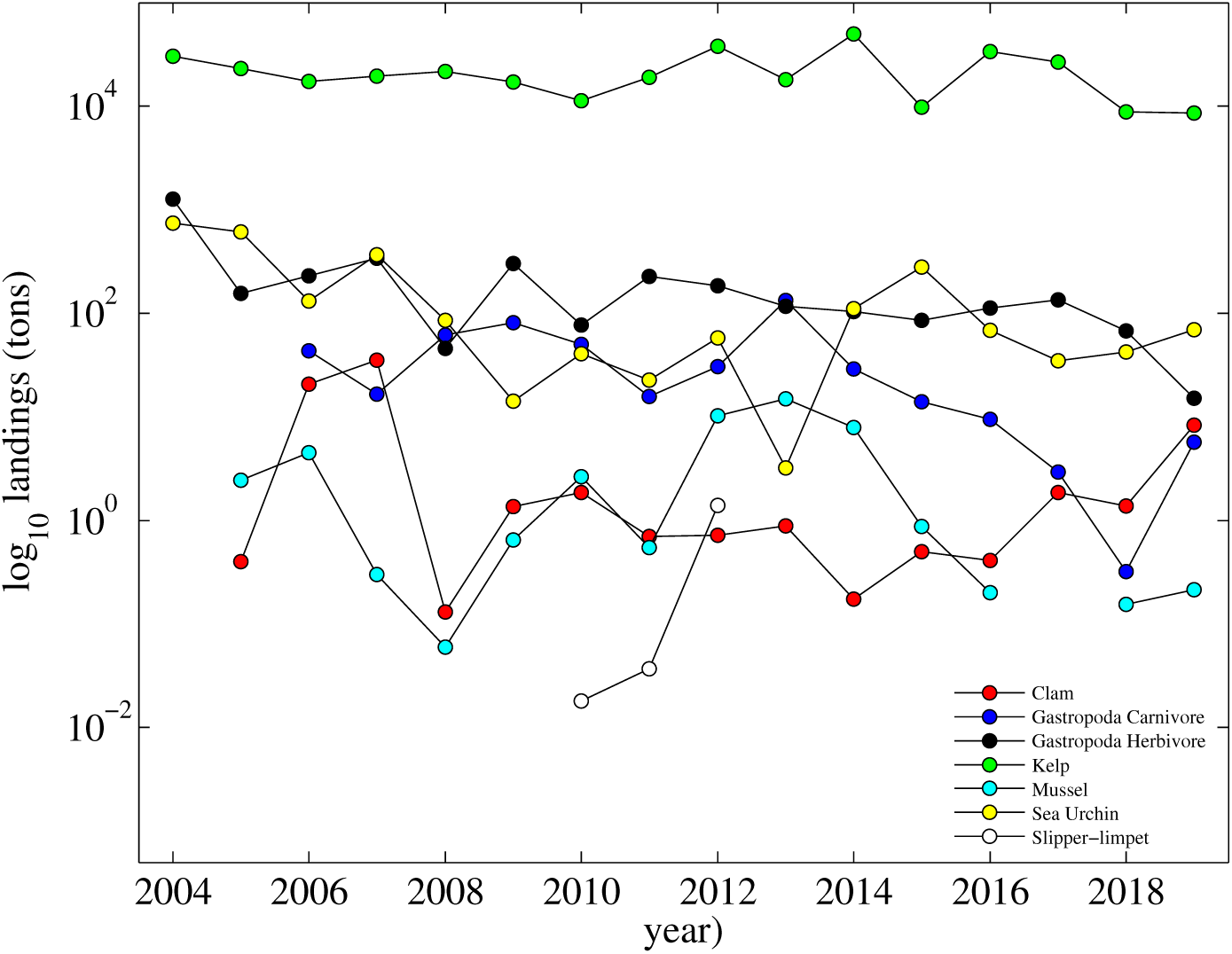
Cumulative annual landings (log_10_(tons)) of the main benthic fisheries reported to IFOP summed across the ports of Paposo and Taltal between 2004 and 2019. Years with no values did not report landings

### 3.2 Temporal–environmental changes in shellfish assemblages

The conditional Deviance Information Criterion, a compromise between fit and complexity, selected the model that used the three SST estimates as a latent variable for the Holocene and the mean for modern assemblages, respectively. On the other hand, all other information criteria indicated that the 50*^th^* SST percentile (mean) provided a more accurate model for the former and the 5*^th^* percentile of SST (min) for the latter. The different information criteria for all possible models are provided in Supplementary Tables 3 and 4. Visual inspection of ordination plots for both latent variables yielded comparable groupings; for simplicity we present ordinations using the mean for both periods. For the Holocene, the ordination of the chronocultural periods using the summed MNI for each functional group and the 50*^th^* percentile of *δ*^18^O SST as a latent variable showed three broadly distinct groupings (Figure 4). The first grouping is associated to soft bottom habitats with bottom-feeding gastropods and slipper limpets and some of the coldest periods of the Holocene (Archaic I and VI). The second grouping is the diverse kelp forest group: sea urchins together with barnacles and decapods, and closely associated to carnivorous and herbivorous gastropods. The former are predatory gastropods, mainly *C. concholepas*, and the latter *Fisurella* spp. keyhole limpets and chitons, respectively (Table 2). These first two grouping are loosely aligned with the colder periods of the Holocene.

**Figure 4:**
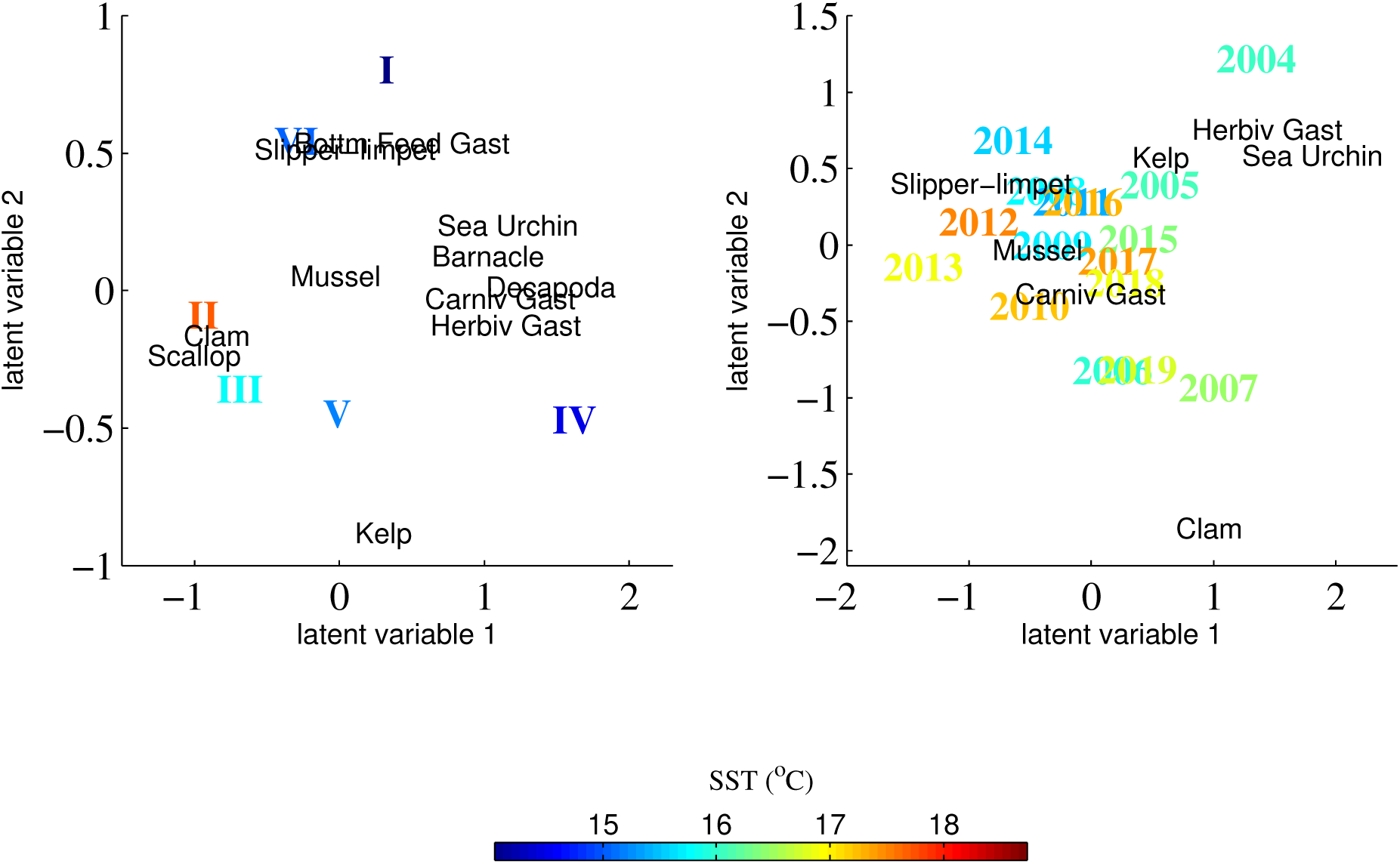
Ordination of the archaeological (left panel) and modern fisheries (right panel) calculated using functional groups and the 50*^th^* percentile of *δ*^18^O SST and MODIS SST values for each cultural period and year, respectively, as a latent variable. The color of the Archaic periods (left panel) and years (right panel) correspond to SST values as indicated by the colorbar at the bottom.

Further in ordination space are less abundant filter-feeding soft bottom bivalves (clams and scallops) tightly clustered around the warmest periods of the Holocene (Archaic II and III). Mussels are isolated in between the groups associated to the warmer and cooler periods, while kelp, indicated by its proxy, the epibiont *S. scurra*, is isolated in the ordination space (Figure 4, left panel). For the 16-year period used for the ordination of modern fisheries, and using the 50*^th^* percentile of SST as a latent variable, we observed a clustering of landings of carnivorous gastropods, chiefly *C. concholepas*, together with other groups that represented a small fraction of landings (i.e. mussels, slipper limpets). This group of species appeared in association with years alternating between the warmest and colder conditions, both el Niño (2010 and 2016-2017) and La Niña (2008 and 2010-2011) years. Kelps, together with sea urchins and herbivorous gastropods grouped closely and in association with cooler years. Clams were separated from any discernible grouping.

## 4 Discussion

### 4.1 Compositional and functional stability in the Atacama coast

Our direct comparison of taxonomic and functional composition between archaeological and ecological data indicates that the benthic ecosystems from the coast of the hyperarid Atacama Desert have not experienced major The compositional similarity extended to the functional level, with herbivorous gastropods and sea urchins, followed by carnivorous gastropods, were among the most common fisheries groups in the past, and remain so in the present (Table 3, Fig. 3). Taken together, the identity of non-shared species between the archaeological and modern set of species was largely a matter of the habitats covered by our sampling design. The lack of biogeographical-scale compositional changes in the past were set against important oceanographic changes in the coastal zone from the early to late Holocene^30,98^. The latitudinal structure of the large-scale SST field shifted during the transition from early to mid-Holocene between southern Peru and northern-central Chile, with intense coastal upwelling displaced equatorward^28,30,50^. However, the composition of the benthic assemblage for the Taltal area (25^◦^S) remained consistent with its biogeographic province within the Humboldt current system, amid two biogeographic breaks around 12^◦^ in southern Perú and 30^◦^S in central Chile. Both are marked by abrupt transitions in the nearshore upwelling regime ^68,87,76^. To our knowledge, only one rocky shore species has recently and permanently expanded its geographic range poleward of 30^◦^S: *S. viridula* ^95,1^ and the archaeological and ecological observations did not record biota occurring equatorward of 12^◦^ from Taltal southward?. The lack of change across the Holocene, including the past decade, stands in stark contrast with the dramatic changes currently taking place worldwide, particularly in kelp forest ecosystems^113,104,13^.

Paleoceanographic data indicates that upwelling-favorable conditions prevailed over much of the Holocene for the arid coastal region along western south America with recent evidence indicating that the intensity of upwelling has increased in the past decades^27,63,55,30,98,117,50^. Thus, elevated benthic primary and secondary production seems bound to remain as a feature of the region in the future.

Covariation in abundance across trophic levels both in the modern and the archaeological assemblage is suggestive of bottom-up forcing of community structure^72,78,118^. We do not have a direct proxy of upwelling intensity or prevalence, yet our results point out that SST dynamics play an important role for the exploited benthic populations both in the archaeological records and in modern fisheries landings. Many of the species that are closely associated to the coldest periods of the Holocene (Archaic I and VI) either lack a pelagic larval stage or lay benthic egg masses (i.e. trochidae, muricidae). Upwelling circulation strongly influence benthic community structure through changes in larval supply for species with pelagic larvae^20,80,76^. On the other hand, the diverse set of species that comprised the kelp forest ecosystem were grouped around similar temperature patterns (Figure 4). The kelp forest group is more clear in the archaeological data owing to its higher functional diversity, yet they are present but not currently fished in our reference region, except for mussels^36^. In the modern association carnivorous gastropods, chiefly *C. concholepas*, despite being separated from the kelp forest group are located together with mussels in ordination space, similar to the archaeological assemblage (Figure 4). Intertidal and subtidal kelp stands are under intense exploitation in the present and their utilization was widespread in the past^5,103,59,25^. The continued exploitation of kelps, and the distinct habitat they define, implies a top-down effect on community structure driven by patterns of human consumption that is altering community composition in benthic environments^33,39,90,77,26^. In the case of soft-bottom habitats, our surveys did not target sandy areas, which are scarce along the Atacama coast^107^. Landings indicate that local artisanal fishers do not exploit extensively soft bottom species, which were common in shell middens. For example, wild populations of Peruvian or Callico scallops, *Argopecten purpuratus*, present in shell midden data at low densities, do not sustain a commercial fishery along the Chilean portion of their geographic range^114^. Archaeological bivalve shells of clams or scallops likely correspond to transient populations established during periods of accumulation of sand over wet periods^110,107,43^. Increased input of sediment, indicative of rainfall and warmer conditions is also associated to warmer waters, weakened upwelling conditions, and increased ENSO activity^86,29,30^. An ENSO-like oceanographic pattern has been repeatedly related to increased larval delivery of *A. purpuratus* in the present^27,114,21^. As such, the association between warmer conditions during the Archaic II and II, and the presence of clams and scallops reinforces the notion of an environmental feedback between benthic community structure and human choices exacerbated by an ENSO-like climatic pattern^22^.

### 4.2 Cultural change, coastal adaptations and climate

Prehistoric societies experienced important cultural transitions in association with environmental change that were key to long-term resilience; different examples around the world suggest a pattern of social reorganization and continuity^99,71,61,42^. In our study region, maritime specialization coincides with the environmental change from early to mid-Holocene, the Archaic II period^92,96^, the warmest ocean conditions in our record^50^. Fishing tools such as shellfish hooks and fishing weights appear for the first time during this cultural period. They become widespread during Archaic III in the Taltal area^5,51,96,92^ and along the northern coast of Chile^18,102^. Technological change during the mid Holocene has been interpreted as a process towards a semi-sedentary settlement system, greater social complexity and a fully specialized maritime economy^96,92^. Under intense upwelling conditions the coastal front may be located hundreds of kilometers offshore; warmer conditions nearshore imply the shoreward displacement of the coastal front and the fauna utilizing the offshore pelagic habitat ^6,15,3^. The increased presence of bone harpoons and offshore fish remains during this period provides indirect evidence of the use of watercraft^85,12,92^, which could have been used to exploit the pelagic assemblage. In the case of our shell midden data, clams and scallop were clearly grouped to these warmer SST conditions (Figure 4), which again suggest that together with maritime specialization, human gathering practices were exploring new habitats with higher natural resource abundances triggered by changing climatic conditions. After 4500 cal yr BP, at the beginning of the late Holocene (Archaic IV), when SST was cooling and closer to its long-term mean (Figure 2A), maritime and coastal subsistence continued to be predominant. However, shell fishhooks were no longer manufactured and habitational stone structures were abandoned along the northern coast of Chile, marking important cultural transformations ^32,96,4^. The cooling SST pattern of this period is associated with the kelp forest assemblage, a more diversified subsistence economy, and changing cultural norms^96^. Cultural and technological changes during the late Holocene also coincide with the onset of the modern ENSO pattern of oceanographic and hydrological variability^29,22^. Warmer water anomalies during El Niño can trigger an increase in occasional but torrential coastal precipitation^119^ and a significant recharge of underground waters^64^. On the other hand, cold water anomalies during La Niña periods are associated with an increased probability of drought^23^. Hence, fostering adaptive capacity to an intensification of ENSO-like climatic variability seems of special importance given current predictions along the eastern Pacific under greenhouse warming^24,115^. The predicted transition of the ENSO climatic pattern, from an irregular regime with moderate amplitude to a highly regular oscillation of intensifying amplitude in a warmer climate ^106^, could be a harbinger of major ecological and cultural changes. The latter is highly relevant as the speed of change forecast towards the second part of the century has never been observed before, testing the adaptive capacity of small-scale fishers and coastal managers in the region^57,38^.

## 5 Conclusion

The socioecological systems of Taltal from where ancient fisher communities derived their sustenance experienced broad changes under fluctuating environmental conditions ^71,101^. The close and sustained relationship around kelp forest ecosystems in the past underlines the need for a cautionary approach to their conservation and management following the unprecedented exploitation of their foundational benthic species in the present^90,59,25,26,77^. Most of the adaptive decisions synthesized in our analyses, although focused on shellfish are convergent with the current guidance for cultivating adaptive capacity to a changing climate in a social ecological system by fostering flexibility, agency, learning, organization organization and assets^38^. Our data and analyzes showed that modern benthic fisheries, which are bounded to their administrative region, but operate in areas of open access, are relatively flexible and able to deploy different assets as they clearly switch between functional groups (Figure 3). However, modern fisheries are bound by a regulatory framework that precludes the deployment of strategic flexibility and the ability to implement new knowledge and organize their agency, for instance, by tracking ongoing changes in fisheries distribution and allocating resources differentially, or changing the species under management^112,58,37,9^. Implementing the lessons of the past may require new approaches in the future. Together, our results have four main conclusions. First, around the latitude of Taltal (25^◦^S), the trans-Holocene archaeological shellfish record does not show major compositional changes across several climatic transitions. No biogeographic transgressions are evident. Secondly, different SST patterns are associated to the distinctive shellfish groupings that have dominated the benthic fishery of the region. Third, the stability of the assemblages gathered and used by people though time highlight the relevance of certain mollusks species and groups such as sea urchins and gastropods in people’s life, independently of the degree of industrialization of the economies. Adaptive capacity should focus around the stewardship of coastal kelp ecosystems as a primary concern^112,57,36^. Finally, we remark on the potential for multidisciplinary studies to shed light on complex patterns of coupled human-environmental changes and provide some clues for the development and fostering of adaptive capacity in coastal socioecological systems.

## 6 Acknowledgements

This work was supported by FONDECYT 1080666, 1110196, 11200953, 1151203,1181300, 1221699 and the Millenium Institute for Coastal Socio-Ecology (SECOS) ICN 015. AC was supported by the Horizon Europe Research and Innovation Programme (grant agreements no 101060072 for the ACTNOW project and no 101213933 for the INSPIRI project); the Pew Marine Fellows Program; and France Filìere P^eche (grant agreement PH/2022/10 for the ADAPT project). Finally, we would like to thank students and colleagues that helped and participated in the several field seasons that were necessary to collect, process and analyze archaeological shellfish data and carry out modern intertidal zone surveys. Our gratitude to Manuel Nuñez for excellent team work while collecting data in the field and on the long drives to Taltal.

## Bibliography

[1] M. A. Aguilera, N. Valdivia, and B. R. Broitman. Spatial niche differentiation and coexistence at the edge: co-occurrence distribution patterns in *Scurria* limpets. Marine Ecology Progress Series, 483:185–198, 2013. doi: 10.3354/meps10293.

[2] A. F. Ainis, R. L. Vellanoweth, Q. G. Lapeña, and C. S. Thornber. Using non-dietary gastropods in coastal shell middens to infer kelp and seagrass harvesting and paleoenvironmental conditions. Journal of Archaeological Science, 49:343–360, 9 2014. doi: 10.1016/J.JAS.2014.05.024.

[3] D. G. Ainley, K. D. Dugger, R. G. Ford, S. D. Pierce, D. C. Reese, R. D. Brodeur, C. T. Tynan, and J. A. Barth. Association of predators and prey at frontal features in the california current: Competition, facilitation, and co-occurrence. Marine Ecology Progress Series, 389:271–294, 9 2009. doi: 10.3354/meps08153.

[4] V. Alcalde and C. Flores. Variabilidad morfológica de anzuelos de concha de *Choromytilus chorus* (8.500-45.00 cal ap), costa sur del desierto de Atacama, Taltal, Chile. Latin American Antiquity, 31:664–682, 2020. doi: 10.1017/laq.2020.29.

[5] V. Alcalde, C. Flores, J. Guardia, L. Olguín, and B. R. Broitman. Marine invertebrates as proxies for early kelp use along the western coast of south america. Frontiers in Earth Science, 11, 2023. doi: 10.3389/feart.2023.1148299.

[6] J. Alheit and M. Niquen. Regime shifts in the humboldt current ecosystem. Progress in Oceanography, 60(2):201–222, 2004. ISSN 0079-6611. doi: 10.1016/j.pocean.2004.02.006.

[7] P. Andrade and D. Salazar. Revisitando Morro Colorado: comparaciones y propuestas preliminares en torno a un conchal arcaico en las costas de Taltal. Taltalia, 4:63–83, 2011.

[8] C. Anguita, S. Gelcich, M. Aldana, and J. Pulgar. Exploring the influence of upwelling on the total allowed catch and harvests of a benthic gastropod managed under a territorial user rights for fisheries regime along the chilean coast. Ocean and Coastal Management, 195, 2020. ISSN 09645691. doi: 10.1016/j.ocecoaman.2020.105256.

[9] N. M. Arias and W. B. Stotz. Management policies impact fishers’ income beyond tolerable limits: The case of the chilean turfs in central chile. Marine Policy, 157: 105834, 11 2023. doi: 10.1016/J.MARPOL.2023.105834.

[10] B. Ballester and F. Gallardo. Prehistoric and historic networks on the Atacama desert coast (northern Chile). Antiquity, 85:875–889, 2011. doi: 10.1017/S0003598X0006837X.

[11] R. T. Barber and F. P. Chavez. Biological consequences of el niño. Science, 222: 1203–1210, 12 1983. doi: 10.1126/SCIENCE.222.4629.1203.

[12] P. Béarez, F. Fuentes-Mucherl, S. Rebolledo, D. Salazar, and L. Olguín. Billfish foraging along the northern coast of Chile during the Middle Holocene (7400–5900cal BP). Journal of Anthropological Archaeology, 41(March):185–195, 2016. doi: 10.1016/j.jaa.2016.01.002.

[13] R. Beas-Luna, F. Micheli, C. B. Woodson, M. Carr, D. Malone, J. Torre, C. Boch, J. E. Caselle, M. Edwards, J. Freiwald, S. L. Hamilton, A. Hernandez, B. Konar, K. J. Kroeker, J. Lorda, G. Montaño-Moctezuma, and G. Torres-Moye. Geographic variation in responses of kelp forest communities of the california current to recent climatic changes. Global Change Biology, 26:6457–6473, 11 2020. ISSN 13652486. doi: 10.1111/gcb.15273.

[14] T. W. Bell, J. G. Allen, K. C. Cavanaugh, and D. A. Siegel. Three decades of variability in California’s giant kelp forests from the Landsat satellites. Remote Sensing of Environment, 238, 2020. doi: 10.1016/j.rse.2018.06.039.

[15] A. Bertrand, J. Habasque, T. Hattab, N. T. Hintzen, R. Oliveros-Ramos, M. Gutiérrez, H. Demarcq, and F. Gerlotto. 3-D habitat suitability of jack mackerel *Trachurus murphyi* in the Southeastern Pacific, a comprehensive study. Progress in Oceanography, 146:199–211, 8 2016. doi: 10.1016/j.pocean.2016.07.002.

[16] L. R. Binford. Butchering, sharing, and the archaeological record. Journal of Anthropological Archaeology, 3(3):235–257, 1984. doi: 10.1016/0278-4165(84)90003-5.

[17] C. A. Blanchette, E. A. Wieters, B. R. Broitman, B. P. Kinlan, and D. R. Schiel. Trophic structure and diversity in rocky intertidal upwelling ecosystems: A comparison of community patterns across California, Chile, South Africa and New Zealand. Progress in Oceanography, 83(1-4):107–116, 2009. doi: 10.1016/j.pocean.2009.07.038.

[18] G. Boisset, A. Llagostera, and E. Salas. Excavaciones arqueológicas en Caleta Abtao, Antofagasta. Actas del V Congreso Nacional de Arqueología, La Serena, pages 75–112, 1969.

[19] B. R. Broitman, S. A. Navarrete, F. Smith, and S. D. Gaines. Geographic variation of southeastern Pacific intertidal communities. Marine Ecology Progress Series, 224: 21–34, 2001. doi: 10.3354/meps224021.

[20] B. R. Broitman, C. A. Blanchette, B. A. Menge, J. Lubchenco, C. Krenz, M. Foley, P. T. Raimondi, D. Lohse, and S. D. Gaines. Spatial and temporal patterns of invertemmubrate recruitment along the West Coast of the United States. Ecological Monographs, 78(3):403–421, 2008. doi: 10.1890/06-1805.1.

[21] B. R. Broitman, C. Lara, R. P. Flores, G. S. Saldías, A. Piñones, A. Pinochet, A. G. Mejía, and S. A. Navarrete. Environmental variability and larval supply to wild and cultured shellfish populations. Aquaculture, 548:737639, 2 2022. doi: 10.1016/j.aquaculture.2021.737639.

[22] J. M. Broughton, B. F. Codding, J. T. Faith, K. A. Mohlenhoff, R. Gruhn, J. Brenner-Coltrain, and I. A. Hart. El Niño frequency threshold controls coastal biotic communities. Science, 377(6611):1202–1205, sep 2022. doi: 10.1126/science.abm1033.

[23] W. Cai, M. J. McPhaden, A. M. Grimm, R. R. Rodrigues, A. S. Taschetto, R. D. Garreaud, B. Dewitte, G. Poveda, Y.-G. Ham, A. Santoso, B. Ng, W. Anderson, G. Wang, T. Geng, H.-S. Jo, J. A. Marengo, L. M. Alves, M. Osman, S. Li, L. Wu, C. Karamperidou, K. Takahashi, and C. Vera. Climate impacts of the El Niño–Southern Oscillation on South America. Nature Reviews Earth and Environment, 1(4):215–231, 2020. doi: 10.1038/s43017-020-0040-3.

[24] W. Cai, A. Santoso, M. Collins, B. Dewitte, C. Karamperidou, J. S. Kug, M. Lengaigne, M. J. McPhaden, M. F. Stuecker, A. S. Taschetto, A. Timmermann, L. Wu, S. W. Yeh, G. Wang, B. Ng, F. Jia, Y. Yang, J. Ying, X. T. Zheng, T. Bayr, J. R. Brown, A. Capotondi, K. M. Cobb, B. Gan, T. Geng, Y. G. Ham, F. F. Jin, H. S. Jo, X. Li, X. Lin, S. McGregor, J. H. Park, K. Stein, K. Yang, L. Zhang, and W. Zhong. Changing el niño–southern oscillation in a warming climate. Nature Reviews Earth and & Environment, 2:628–644, 9 2021. ISSN 2662138X. doi: 10.1038/s43017-021-00199-z.

[25] P. Carbajal, A. Gamarra Salazar, P. J. Moore, and A. Pérez-Matus. Different kelp species support unique macroinvertebrate assemblages, suggesting the potential community-wide impacts of kelp harvesting along the Humboldt Current System. Aquatic Conservation: Marine and Freshwater Ecosystems, 32(1):14–27, dec 2022. doi: 10.1002/aqc.3745.

[26] D. M. Carranza, G. C. Stotz, J. A. Vásquez, and W. B. Stotz. Trends in the effects of kelp removal on kelp populations, herbivores, and understory algae. Global Ecology and Conservation, 49:e02805, 2024. doi: 10.1016/j.gecco.2024.e02805.

[27] M. Carré, I. Bentaleb, D. Blamart, N. Ogle, F. Cardenas, S. Zevallos, R. M. Kalin, L. Ortlieb, and M. Fontugne. Stable isotopes and sclerochronology of the bivalve *Mesodesma donacium*: Potential application to Peruvian paleoceanographic reconstructions. Palaeogeography, Palaeoclimatology, Palaeoecology, 228(1-2):4–25, 2005. doi: 10.1016/j.palaeo.2005.03.045.

[28] M. Carré, M. Azzoug, I. Bentaleb, B. M. Chase, M. Fontugne, D. Jackson, M. P. Ledru, A. Maldonado, J. P. Sachs, and A. J. Schauer. Mid-Holocene mean climate in the south eastern Pacific and its influence on South America. Quaternary International, 253:55–66, mar 2012. doi: 10.1016/j.quaint.2011.02.004.

[29] M. Carré, J. P. Sachs, S. Purca, A. J. Schauer, P. Braconnot, R. A. Falcón, M. Julien, and D. Lavallée. Holocene history of ENSO variance and asymmetry in the eastern tropical Pacific. Science, 345(6200):1045–1048, aug 2014. doi: 10.1126/science.1252220.

[30] M. Carré, D. Jackson, A. Maldonado, B. M. Chase, and J. P. Sachs. Variability of ^14^C reservoir age and air-sea flux of CO_2_ in the Peru-Chile upwelling region during the past 12,000 years. Quaternary Research (United States), 85(1):87–93, 2016. doi: 10.1016/j.yqres.2015.12.002.

[31] R. W. Casteel. Some archaeological uses of fish remains. American Antiquity, 37(3): 404–419, 1972. doi: 10.2307/278439.

[32] J. Castelleti. Patrón de asentamiento y uso de recursos a través de la secuencia ocupacional prehispana en la costa de Taltal. Master’s thesis, Universidad Católica del Norte, 2007.

[33] J. C. Castilla. Coastal marine communities: Trends and perspectives from human-exclusion experiments. Trends in Ecology and Evolution, 14:280–282, 7 1999. doi: 10.1016/S0169-5347(99)01602-X.

[34] J. C. Castilla and P. Camus. The Humboldt-el Niño scenario:coastal benthic resources and anthropogenic influences, with particular reference to the 1982/1983 ENSO. South African Journal of Marine Sciences, 12:703–712, 1992.

[35] F. P. Chavez and M. Messié. A comparison of Eastern Boundary Upwelling Ecosystems. Progress in Oceanography, 83(1-4):80–96, jan 2009. doi: 10.1016/j.pocean.2009.07.032.

[36] A. Chevallier, B. R. Broitman, N. Barahona, C. Vicencio-Estay, F. K. Hui, P. Inchausti, and W. B. Stotz. Diversity of small-scale fisheries in Chile: Environmental patterns and biogeography can inform fisheries management. Environmental Science and Policy, 124(December 2020):33–44, 2021. doi: 10.1016/j.envsci.2021.06.002.

[37] A. Chevallier, W. Stotz, M. Ramos, and J. Mendo. The Humboldt Current Large Marine Ecosystem (HCLME), a challenging scenario for modelers and their contribution for the manager, pages 27–51. Springer International Publishing, 2021.

[38] J. E. Cinner, W. N. Adger, E. H. Allison, M. L. Barnes, K. Brown, P. J. Cohen, S. Gelcich, C. C. Hicks, T. P. Hughes, J. Lau, N. A. Marshall, and T. H. Morrison. Building adaptive capacity to climate change in tropical coastal communities. Nature Climate Change, 8:117–123, 2018. doi: 10.1038/s41558-017-0065-x.

[39] F. C. Coleman and S. L. Williams. Overexploiting marine ecosystem engineers: potential consequences for biodiversity. Trends in Ecology and Evolution, 17(1):40–44, 2002. doi: 10.1016/S0169-5347(01)02330-8.

[40] B. I. Crona, T. V. Holt, M. Petersson, T. M. Daw, and E. Buchary. Using social–ecological syndromes to understand impacts of international seafood trade on small-scale fisheries. Global Environmental Change, 35:162–175, 11 2015. doi: 10.1016/J.GLOENVCHA.2015.07.006.

[41] P. K. Dayton, M. J. Tegner, P. B. Edwards, and K. L. Riser. Temporal and spatial scales of kelp demography: The role of oceanographic climate. Ecological Monographs, 69(2):219–250, 1999. doi: 10.1890/0012-9615(1999)069[0219:TASSOK]2.0.CO;2.

[42] J. G. de Souza, M. Robinson, S. Y. Maezumi, J. Capriles, J. A. Hoggarth, U. Lombardo, V. F. Novello, J. Apáestegui, B. Whitney, D. Urrego, D. T. Alves, S. Rostain, M. J. Power, F. E. Mayle, F. W. da Cruz, H. Hooghiemstra, and J. Iriarte. Climate change and cultural resilience in late pre-columbian amazonia. Nature Ecology and Evolution, 3:1007–1017, 7 2019. ISSN 2397334X. doi: 10.1038/S41559-019-0924-0;SUBJMETA.

[43] J. L. Diederich, V. Wennrich, R. Bao, C. Büttner, A. Bolten, D. Brill, S. Buske, E. Campos, E. Fernández-Galego, P. Gödickmeier, L. Ninnemann, M. Reyers, B. Ritter, L. Ritterbach, C. Rolf, S. Scheidt, T. J. Dunai, and M. Melles. A 68 ka precipitation record from the hyperarid core of the Atacama Desert in northern Chile. Global and Planetary Change, 184:103054, 2020. doi: 10.1016/j.gloplacha.2019.103054.

[44] T. D. Dillehay, C. Ramírez, M. Pino, M. B. Collins, J. Rossen, and J. D. Pino-Navarro. Monte verde: Seaweed, food, medicine, and the peopling of south america. Science, 320:784–786, 5 2008. doi: 10.1126/science.1156533.

[45] J. M. Erlandson, M. H. Graham, B. J. Bourque, D. Corbett, J. A. Estes, and R. S. Steneck. The kelp highway hypothesis: marine ecology, the coastal migration theory, and the peopling of the americas. The Journal of Island and Coastal Archaeology, 2 (2):161–174, 2007.

[46] R. Escribano, G. Daneri, L. Farías, V. A. Gallardo, H. E. González, D. Gutiérrez, C. B. Lange, C. E. Morales, O. Pizarro, O. Ulloa, and M. Braun. Biological and chemical consequences of the 1997-1998 el niño in the chilean coastal upwelling system: A synthesis. Deep-Sea Research Part II: Topical Studies in Oceanography, 51:2389–2411, 10 2004. ISSN 09670645. doi: 10.1016/j.dsr2.2004.08.011.

[47] J. A. Estes and G. J. Vermeij. History’s legacy: Why future progress in ecology demands a view of the past. Ecology, 103:e3788, 11 2022. doi: 10.1002/ECY.3788.

[48] P. B. Fenberg, B. A. Menge, P. T. Raimondi, and M. M. Rivadeneira. Biogeographic structure of the northeastern Pacific rocky intertidal: The role of upwelling and dispersal to drive patterns. Ecography, 38(1):83–95, 2015. doi: 10.1111/ecog.00880.

[49] C. Flores. Importance of small-scale paleo-oceanographic conditions to interpret changes in size of California mussel (*Mytilus californianus*). Late Holocene, Santa Cruz island, California. Quaternary International, 427:137–150, 2017. ISSN 1040-6182. doi: 10.1016/j.quaint.2016.01.036.

[50] C. Flores and B. R. Broitman. Nearshore paleoceanogaphic conditions through the holocene: Shell carbonate from archaeological sites of the Atacama desert coast. Palaeogeography, Palaeoclimatology, Palaeoecology, 562:110090, 2021. doi: 10.1016/j.palaeo.2020.110090.

[51] C. Flores, V. Figueroa, and D. Salazar. Middle Holocene production of mussel shell fishing artifacts on the coast of Taltal (25^◦^S), Atacama desert, Chile. The Journal of Island and Coastal Archaeology, 11:411–424, 2016. doi: 10.1080/15564894.2015.1105884.

[52] C. Flores, E. M. Gayo, D. Salazar, and B. R. Broitman. *δ*^18^O of Fissurella maxima as a proxy for reconstructing Early Holocene sea surface temperatures in the coastal Atacama desert (25^◦^S). Palaeogeography, Palaeoclimatology, Palaeoecology, 499:22–34, 2018. doi: 10.1016/j.palaeo.2018.03.031.

[53] M. Fontugne, M. Carré, I. Bentaleb, M. Julien, and D. Lavallée. Radiocarbon reservoir age variations in the south Peruvian upwelling during the Holocene. Radiocarbon, 46 (2):531–537, 2004. doi: 10.1017/S003382220003558X.

[54] E. Fragkopoulou, E. A. Serrão, O. De Clerck, M. J. Costello, M. B. Araújo, C. M. Duarte, D. Krause-Jensen, and J. Assis. Global biodiversity patterns of marine forests of brown macroalgae. Global Ecology and Biogeography, 31(4):636–648, apr 2022. doi: 10.1111/geb.13450.

[55] M. García-Reyes, W. J. Sydeman, D. S. Schoeman, R. R. Rykaczewski, B. a. Black, A. J. Smit, and S. J. Bograd. Under pressure: Climate change, upwelling, and eastern boundary upwelling ecosystems. Frontiers in Marine Science, 2:1–10, 2015. doi: 10.3389/fmars.2015.00109.

[56] GEBCO Compilation Group. General Bathymetric Chart of the Oceans, 2023. GEBCO 2023 Grid, 10.5285/f98b053b-0cbc-6c23-e053-6c86abc0af7b.

[57] S. Gelcich, T. P. Hughes, P. Olsson, C. Folke, O. Defeo, M. Fernández, S. Foale, L. H. Gunderson, C. Rodríguez-Sickert, M. Scheffer, R. S. Steneck, and J. C. Castilla. Navigating transformations in governance of chilean marine coastal resources. Proceedings of the National Academy of Sciences of the United States of America, 107:16794–16799, 9 2010. doi: 10.1073/pnas.1012021107.

[58] J. E. González, B. Yannicelli, and W. Stotz. The interplay of natural variability, productivity and management of the benthic ecosystem in the Humboldt Current System: Twenty years of assessment of *Concholepas concholepas* fishery under a TURF management system. Ocean and Coastal Management, 208:105628, 7 2021. doi: 10.1016/j.ocecoaman.2021.105628.

[59] A. Gouraguine, P. Moore, M. T. Burrows, E. Velasco, L. Ariz, L. Figueroa-Fábrega, R. Muñoz-Cordovez, I. Fernandez-Cisternas, D. Smale, and A. Pérez-Matus. The intensity of kelp harvesting shapes the population structure of the foundation species *Lessonia trabeculata* along the chilean coastline. Marine Biology, 168:1–9, 5 2021. doi: 10.1007/S00227-021-03870-7.

[60] D. K. Grayson. Quantitative Zooarchaeology: Topics in the Analysis of Archaelogical Faunas. Elsevier, 1984.

[61] A. S. Groesbeck, K. Rowell, D. Lepofsky, and A. K. Salomon. Ancient clam gardens increased shellfish production: Adaptive strategies from the past can inform food security today. PLOS ONE, 9:e91235, 3 2014. doi: 10.1371/JOURNAL.PONE.0091235.

[62] J. Guardia. El recurso malacológico y su consumo alimenticio en los grupos cazadores recolectores Pescadores en la costa de Taltal-Paposo (12.000-1.500 años AP) Memoria para optar al título de Arqueóloga, Universidad SEK, 2020.

[63] D. Gutiérrez, I. Bouloubassi, A. Sifeddine, S. Purca, K. Goubanova, M. Graco, D. Field, L. Méjanelle, F. Velazco, A. Lorre, R. Salvatteci, D. Quispe, G. Vargas, B. Dewitte, and L. Ortlieb. Coastal cooling and increased productivity in the main upwelling zone off Peru since the mid-twentieth century. Geophysical Research Letters, 38:L07603, 4 2011. doi: 10.1029/2010GL046324.

[64] C. Herrera and E. Custodio. Origen de las aguas de pequeños manantiales de la Costa del norte de Chile, en las cercanías de Antofagasta. Andean Geology, 41(2):314–341, 2014. doi: 10.5027/andgeoV41n2-a03.

[65] F. K. Hui. BORAL: Bayesian ordination and regression analysis. R package version 1.9, 2020.

[66] F. K. Hui, S. Taskinen, S. Pledger, S. D. Foster, and D. I. Warton. Model-based approaches to unconstrained ordination. Methods in Ecology and Evolution, 6(4):399–411, apr 2015. doi: 10.1111/2041-210X.12236.

[67] N. P. Jew, S. M. Fitzpatrick, and K. J. Sullivan. *δ*^18^O analysis of *Donax denticulatus*: Evaluating a proxy for sea surface temperature and nearshore paleoenvironmental reconstructions for the northern Caribbean. Journal of Archaeological Science: Reports, 8:216–223, 2016. doi: 10.1016/j.jasrep.2016.06.018.

[68] C. Lara, G. S. Saldías, B. Cazelles, M. M. Rivadeneira, P. A. Haye, and B. R. Broitman. Coastal biophysical processes and the biogeography of rocky intertidal species along the south-eastern Pacific. Journal of Biogeography, 46(2):420–431, feb 2019. doi: 10.1111/jbi.13492.

[69] C. Latorre, R. De Pol-Holz, C. Carter, and C. M. Santoro. Using archaeological shell middens as a proxy for past local coastal upwelling in northern Chile. Quaternary International, 427(DECEMBER):128–136, 2017. doi: 10.1016/j.quaint.2015.11.079.

[70] A. Llagostera, G. Weisner, M. Castillo, and M. Cervelino. El complejo Huentelauquén bajo una perspectiva macro-espacial y multidisciplinaria. Contribuciones Arqueológicas, 5:461–480, 2000.

[71] P. A. Marquet, C. M. Santoro, C. Latorre, V. G. Standen, S. R. Abades, M. M. Rivadeneira, B. Arriaza, and M. E. Hochberg. Emergence of social complexity among coastal hunter-gatherers in the Atacama Desert of northern Chile. Proceedings of the National Academy of Sciences of the United States of America, 109(37):14754–14760, aug 2012. doi: 10.1073/pnas.1116724109.

[72] B. A. Menge, J. Lubchenco, M. E. S. Bracken, F. Chan, M. M. Foley, T. L. Freidenburg, S. D. Gaines, G. Hudson, C. Krenz, H. Leslie, D. N. L. Menge, R. Russell, and M. S. Webster. Coastal oceanography sets the pace of rocky intertidal community dynamics. Proceedings of the National Academy of Sciences of the United States of America, 100 (21):12229–12234, oct 2003. doi: 10.1073/pnas.1534875100.

[73] M. Mohtadi, O. E. Romero, and D. Hebbeln. Changing marine productivity off northern Chile during the past 19 000 years: A multivariable approach. Journal of Quaternary Science, 19(4):347–360, 2004. doi: 10.1002/jqs.832.

[74] J. V. Moreno-Mayar, L. Vinner, P. D. B. Damgaard, C. D. L. Fuente, J. Chan, J. P. Spence, M. E. Allentoft, T. Vimala, F. Racimo, T. Pinotti, S. Rasmussen, A. Margaryan, M. I. Orbegozo, D. Mylopotamitaki, M. Wooller, C. Bataille, L. Becerra-Valdivia, D. Chivall, D. Comeskey, T. Devìese, D. K. Grayson, L. George, H. Harry, V. Alexandersen, C. Primeau, J. Erlandson, C. Rodrigues-Carvalho, S. Reis, M. Q. Bastos, J. Cybulski, C. Vullo, F. Morello, M. Vilar, S. Wells, K. Gregersen, K. L. Hansen, N. Lynnerup, M. M. Lahr, K. Kjær, A. Strauss, M. Alfonso-Durruty, A. Salas, H. Schroeder, T. Higham, R. S. Malhi, J. T. Rasic, L. Souza, F. R. Santos, A. S. Malaspinas, M. Sikora, R. Nielsen, Y. S. Song, D. J. Meltzer, and E. Willerslev. Early human dispersals within the americas. Science, 362, 12 2018. ISSN 10959203. doi: 10.1126/science.aav2621.

[75] S. A. Navarrete, E. A. Wieters, B. R. Broitman, and J. C. Castilla. Scales of benthic-pelagic coupling and the intensity of species interactions: from recruitment limitation to top-down control. Proceedings of the National Academy of Sciences of the United States of America, 102(50):18046–18051, 2005. doi: 10.1073/pnas.0509119102.

[76] S. A. Navarrete, M. Barahona, N. Weidberg, and B. R. Broitman. Climate change in the coastal ocean: Shifts in pelagic productivity and regionally diverging dynamics of coastal ecosystems. Proceedings of the Royal Society B: Biological Sciences, 289:11–13, 2022. doi: 10.1098/rspb.2021.2772.

[77] S. A. Navarrete, M. I. Ávila Thieme, D. Valencia, A. Génin, and S. Gelcich. Monitoring the fabric of nature: using allometric trophic network models and observations to assess policy effects on biodiversity. Philosophical Transactions of the Royal Society B, 378, 7 2023. doi: 10.1098/RSTB.2022.0189.

[78] K. J. Nielsen and S. A. Navarrete. Mesoscale regulation comes from the bottom-up: Intertidal interactions between consumers and upwelling. Ecology Letters, 7(1):31–41, jan 2004. doi: 10.1046/j.1461-0248.2003.00542.x.

[79] L. Núñez and C. M. Santoro. El tránsito arcaico-formativo en la circumpuna y valles occidentales del centro sur andino: hacia los cambios” neolíticos”. *Chungara*, Revista de Antropología Chilena, 43:487–530, 2011. doi: 10.4067/S0717-73562011000300010.

[80] D. K. Okamoto, S. C. Schroeter, and D. C. Reed. Effects of ocean climate on spatiotemporal variation in sea urchin settlement and recruitment. Limnology and Oceanography, 65:2076–2091, 2020. doi: 10.1002/lno.11440.

[81] L. Olguín, V. Castro, M. P. Castro Torres, I. Peña Villalobos, J. Ruz, and B. Santander. Exploitation of faunal resources by marine hunter-gatherer groups during the Middle Holocene at the Copaca 1 site, Atacama Desert coast. Quaternary International, 373 (April):4–16, 2015. doi: 10.1016/j.quaint.2015.02.004.

[82] L. Olguín, C. Flores, and D. Salazar. Aprovechamiento humano de moluscos marinos en conchales arqueológicos del Holoceno Temprano y Medio (12.000–5.000 años cal A.P.). Costa meridional del Desierto de Atacama. Chile. In H. Hammond and Z. MA, editors, Arqueomalacología: abordajes metodológicos y casos de estudio en el Cono Sur, page 3–34. Fundación de Historia Natural Félix de Azara, Buenos Aires, 2015.

[83] L. Olguín. Aproximaciones a la historia de la actividad minero-metalúrgica indígena en la costa desértica de la Región de Antofagasta: Localidades de Taltal y Paposo. Technical report, Informe arqueo-malacológico. FONDECYT 1080666, 2009–2010.

[84] L. Olguín. Cazadores-recolectores, pescadores y mineros del período Arcaico en la costa de Taltal. Technical report, Informe arqueo-malacológico. FONDECYT 1110196, 2012–2014.

[85] L. Olguín. Trayectoria histórica, cambios ambientales y eventos catastróficos durante el período Arcaico en la costa de Taltal, Norte de Chile. Technical report, Informe arqueo-malacológico. FONDECYT 1151203, 2015.

[86] L. Ortlieb, G. Vargas, and J.-F. Salìege. Marine radiocarbon reservoir effect along the northern Chile–southern Peru coast (14–24^◦^S) throughout the Holocene. Quaternary Research, 75(1):91–103, 2011. doi: 10.1016/j.yqres.2010.07.018.

[87] L. Peluso, B. R. Broitman, M. A. Lardies, R. F. Nespolo, and P. Saenz. Comparative population genetics of congeneric limpets across a biogeographic transition zone reveals common patterns of genetic structure and demographic history. Molecular Ecology, 32 (April):1–14, 2023. doi: 10.1111/mec.16978.

[88] M. L. Pinsky, H. Hillebrand, J. M. Chase, L. H. Antão, M. R. Hirt, U. Brose, M. T. Burrows, B. Gauzens, B. Rosenbaum, and S. A. Blowes. Warming and cooling catalyse widespread temporal turnover in biodiversity. Nature, pages 1–5, 1 2025. ISSN 1476-4687. doi: 10.1038/s41586-024-08456-z. URL https://www.nature.com/articles/s41586-024-08456-z.

[89] X. Power and D. Salazar. Estudio intrasitio de un yacimiento arcaico con arquitectura en la costa de Taltal, Desierto de Atacama, Norte de Chile. Chungara. Revista de Antropología Chilena, 52:183–207, 2020. ISSN 0717-7356. doi: 10.4067/S0717-73562020005001201.

[90] A. Pérez-Matus, A. Ospina-Alvarez, P. A. Camus, S. A. Carrasco, M. Fernandez, S. Gelcich, N. Godoy, F. P. Ojeda, L. M. Pardo, N. Rozbaczylo, M. D. Subida, M. Thiel, E. A. Wieters, and S. A. Navarrete. Temperate rocky subtidal reef community reveals human impacts across the entire food web. Marine Ecology Progress Series, 567:1–16, 3 2017. doi: 10.3354/meps12057.

[91] M. Raghavan, M. Steinrücken, K. Harris, S. Schiffels, S. Rasmussen, M. DeGiorgio, A. Albrechtsen, C. Valdiosera, M. C. Ávila Arcos, A. S. Malaspinas, A. Eriksson, I. Moltke, M. Metspalu, J. R. Homburger, J. Wall, O. E. Cornejo, J. V. Moreno-Mayar, T. S. Korneliussen, T. Pierre, M. Rasmussen, P. F. Campos, P. D. B. Damgaard, M. E. Allentoft, J. Lindo, E. Metspalu, R. Rodríguez-Varela, J. Mansilla, C. Henrickson, A. Seguin-Orlando, H. Malmstöm, T. Stafford, S. S. Shringarpure, A. Moreno-Estrada, M. Karmin, K. Tambets, A. Bergström, Y. Xue, V. Warmuth, A. D. Friend, J. Singarayer, P. Valdes, F. Balloux, I. Leboreiro, J. L. Vera, H. Rangel-Villalobos, D. Pettener, D. Luiselli, L. G. Davis, E. Heyer, C. P. Zollikofer, M. S. P. D. León, C. I. Smith, V. Grimes, K. A. Pike, M. Deal, B. T. Fuller, B. Arriaza, V. Standen, M. F. Luz, F. Ricaut, N. Guidon, L. Osipova, M. I. Voevoda, O. L. Posukh, O. Balanovsky, M. Lavryashina, Y. Bogunov, E. Khusnutdinova, M. Gubina, E. Balanovska, S. Fedorova, S. Litvinov, B. Malyarchuk, M. Derenko, M. J. Mosher, D. Archer, J. Cybulski, B. Petzelt, J. Mitchell, R. Worl, P. J. Norman, P. Parham, B. M. Kemp, T. Kivisild, C. Tyler-Smith, M. S. Sandhu, M. Crawford, R. Villems, D. G. Smith, M. R. Waters, T. Goebel, J. R. Johnson, R. S. Malhi, M. Jakobsson, D. J. Meltzer, A. Manica, R. Durbin, C. D. Bustamante, Y. S. Song, R. Nielsen, and E. Willerslev. Genomic evidence for the pleistocene and recent population history of native americans. Science, 349, 8 2015. doi: 10.1126/SCIENCE.AAB3884.

[92] S. Rebolledo, P. Béarez, D. Salazar, and F. Fuentes. Maritime fishing during the Middle Holocene in the hyperarid coast of the Atacama Desert. Quaternary International, 391: 3–11, 2016. doi: 10.1016/j.quaint.2015.09.051.

[93] E. J. Reitz and E. S. Wing. Zooarchaeology. Cambridge University Press, 1999.

[94] T. C. Rick. Shell midden archaeology: Current trends and future directions. Journal of Archaeological Research 2023 32:3, 32:309–366, 9 2023. doi: 10.1007/S10814-023-09189-9.

[95] M. M. Rivadeneira and M. Fernández. Shifts in southern endpoints of distribution in rocky intertidal species along the south-eastern Pacific coast. Journal of Biogeography, 32(2):203–209, 2005. doi: 10.1111/j.1365-2699.2004.01133.x.

[96] D. Salazar, V. Figueroa, P. Andrade, H. Salinas, L. Olguín, X. Power, S. Rebolledo, S. Parra, H. Orellana, and J. Urrea. Cronología y organización económica de las poblaciones arcaicas de la costa de Taltal. Estudios Atacameños, 50:07–46, 2015. doi: 10.4067/S0718-10432015000100002.

[97] D. Salazar, C. Arenas, P. Andrade, L. Olguín, J. Torres, C. Flores, G. Vargas, S. Rebolledo, C. Borie, and C. Sandoval. From the use of space to territorialisation during the Early Holocene in Taltal, coastal Atacama Desert, Chile. Quaternary International, 473:225–241, 2018. doi: 10.1016/j.quaint.2017.09.035.

[98] R. Salvatteci, R. R. Schneider, T. Blanz, and E. Mollier-Vogel. Deglacial to holocene ocean temperatures in the humboldt current system as indicated by alkenone paleothermometry. Geophysical Research Letters, 46:281–292, 1 2019. doi: 10.1029/2018GL080634.

[99] D. H. Sandweiss, K. A. Maasch, R. L. Burger, J. B. I. Richardson, H. B. Rollins, and A. Clement. Variation in Holocene El Niño frequencies: Climate records and cultural consequences in ancient Peru. Geology, 29(7):603–606, 2001. doi: 10.1130/0091-7613(2001)029⟨0603:VIHENO⟩2.0.CO;2.

[100] C. M. Santoro, J. M. Capriles, E. M. Gayo, M. E. De Porras, A. Maldonado, V. G. Standen, C. Latorre, V. Castro, D. Angelo, and V. McRostie. Continuities and discontinuities in the socio-environmental systems of the Atacama Desert during the last 13,000 years. Journal of Anthropological Archaeology, 46:28–39, 2017. doi: 10.1016/j.jaa.2016.08.006.

[101] C. M. Santoro, E. M. Gayo, C. Carter, V. G. Standen, V. Castro, D. Valenzuela, R. D. Pol-Holz, P. A. Marquet, and C. Latorre. Loco or no loco? Holocene climatic fluctuations, human demography, and community based management of coastal resources in Northern Chile. Frontiers in Earth Science, 5, 2017. doi: 10.3389/feart.2017.00077.

[102] V. Schiappacasse and H. Niemeyer. *Descripción y análisis interpretativo de un sitio arcaico temprano en la Quebrada de Camarones*. Publicación Ocasional 41. Museo Nacional de Historia Natural y Universidad de Tarapacá, 1984.

[103] L. Sitzia, X. Power, D. Zurro, J. P. Maalouf, J. Cárcamo, K. Chandía, J. M. A. Vega, C. Borie, C. Roa, C. Silva, D. Salazar, S. Vivanco, V. Hernández, C. Aliste, S. Ibacache, and R. Lorca. Tracking kelp-type seaweed fuel in the archaeological record through Raman spectroscopy of charred particles: examples from the Atacama Desert coast. Archaeological and Anthropological Sciences, 15:1–31, 11 2023. doi: 10.1007/S12520-023-01860-Y.

[104] S. M. Smith, H. A. Malcolm, E. M. Marzinelli, A. L. Schultz, P. D. Steinberg, and A. Vergés. Tropicalization and kelp loss shift trophic composition and lead to more winners than losers in fish communities. Global Change Biology, 27:2537–2548, 6 2021. doi: 10.1111/gcb.15592.

[105] P. Strub, J. Mesías, V. Montecino, J. Rutlant, and S. Salinas. Coastal ocean circulation off western South America. The sea, Vol 11, 11:273–314, 1998.

[106] M. F. Stuecker, S. Zhao, A. Timmermann, R. Ghosh, T. Semmler, S. S. Lee, J. Y. Moon, F. F. Jin, and T. Jung. Global climate mode resonance due to rapidly intensifying el niño-southern oscillation. Nature Communications, 16:1–13, 10 2025. doi: 10.1038/s41467-025-64619-0.

[107] J.-B. W. Stuut, S. Kasten, F. Lamy, and D. Hebbeln. Sources and modes of terrigenous sediment input to the Chilean continental slope. Quaternary International, 161(1):67–76, 2007. doi: 10.1016/j.quaint.2006.10.041.

[108] M. Thiel, E. C. Macaya, E. Acuña, W. E. Arntz, H. Bastias, K. Brokordt, P. A. Camus, J. C. Castilla, L. R. Castro, M. Cortés, C. P. Dumont, R. Escribano, M. Fernández, J. A. Gajardo, C. F. Gaymer, I. Gomez, A. E. González, H. E. González, P. A. Haye, J.-E. Illanes, J. L. Iriarte, D. A. Lancellotti, G. Luna-Jorquera, C. Luxoro, P. H. Manríquez, V. Marín, P. Muñoz, S. A. Navarrete, E. Perez, E. Poulin, J. Sellanes, H. H. Sepúlveda, W. Stotz, F. Tala, A. Thomas, C. A. Vargas, J. A. Vasquez, and J. A. Vega. The humboldt current system of northern and central chile. oceanographic processes, ecological interactions and socioeconomic feedback. Oceanography and Marine Biology: An Annual Review, 45:195–344, 2007.

[109] O. Ulloa, R. Escribano, S. Hormazabal, R. A. Quinones, R. R. González, and M. Ramos. Evolution and biological effects of the 1997–98 el nino in the upwelling ecosystem off northern chile. Geophysical Research Letters, 28:1591–1594, 2001.

[110] G. Vargas, J. Rutllant, and L. Ortlieb. ENSO tropical-extratropical climate teleconnections and mechanisms for Holocene debris flows along the hyperarid coast of western South America (17°-24°S). Earth and Planetary Science Letters, 249(3-4):467–483, 2006. doi: 10.1016/j.epsl.2006.07.022.

[111] L. Vasquez. Trayectoria histórica, cambios ambientales y eventos catastróficos durante el período arcaico en la costa de Taltal, Norte de Chile. Technical report, Práctica análisis arqueo-malacológico. FONDECYT 1151203, 2015.

[112] J. M. A. Vega, B. R. Broitman, and J. A. Vásquez. Monitoring the sustainability of lessonia nigrescens (laminariales, phaeophyceae) in northern chile under strong harvest pressure. Journal of Applied Phycology, 26:791–801, 2014. doi: 10.1007/s10811-013-0167-4.

[113] A. Vergés, P. D. Steinberg, M. E. Hay, A. G. Poore, A. H. Campbell, E. Ballesteros, K. L. Heck, D. J. Booth, M. A. Coleman, D. A. Feary, W. Figueira, T. Langlois, E. M. Marzinelli, T. Mizerek, P. J. Mumby, Y. Nakamura, M. Roughan, E. van Sebille, A. S. Gupta, D. A. Smale, F. Tomas, T. Wernberg, and S. K. Wilson. The tropicalization of temperate marine ecosystems: Climate-mediated changes in herbivory and community phase shifts. Proceedings of the Royal Society B: Biological Sciences, 281, 7 2014. doi: 10.1098/rspb.2014.0846.

[114] E. von Brand, A. Abarca, G. E. Merino, and W. Stotz. Scallop Fishery and Aquaculture in Chile: A History of Developments and Declines. Developments in Aquaculture and Fisheries Science, 40:1047–1072, 2016. doi: 10.1016/B978-0-444-62710-0.00026-2.

[115] B. Wang, W. Sun, C. Jin, X. Luo, Y.-M. Yang, T. Li, B. Xiang, M. J. McPhaden, M. A. Cane, F. Jin, F. Liu, and J. Liu. Understanding the recent increase in multiyear La Niñas. Nature Climate Change, 13:1075–1081, 2023. doi: 10.1038/s41558-023-01801-6.

[116] D. I. Warton, F. G. Blanchet, R. B. O’Hara, O. Ovaskainen, S. Taskinen, S. C. Walker, and F. K. Hui. So Many Variables: Joint Modeling in Community Ecology. Trends in Ecology and Evolution, 30(12):766–779, dec 2015. doi: 10.1016/j.tree.2015.09.007.

[117] N. Weidberg, A. Ospina-Alvarez, J. Bonicelli, M. Barahona, C. M. Aiken, B. R. Broitman, and S. A. Navarrete. Spatial shifts in productivity of the coastal ocean over the past two decades induced by migration of the Pacific Anticyclone and Bakun’s effect in the Humboldt Upwelling Ecosystem. Global and Planetary Change, 193:103259, 10 2020. doi: 10.1016/j.gloplacha.2020.103259.

[118] E. A. Wieters, B. R. Broitman, and G. M. Branch. Benthic community structure and spatiotemporal thermal regimes in two upwelling ecosystems: Comparisons between South Africa and Chile. Limnology and Oceanography, 54(4):1060–1072, 2009. doi: 10.4319/lo.2009.54.4.1060.

[119] A. Williams, C. M. Santoro, M. A. Smith, and C. Latorre. The impact of ENSO in the Atacama desert and Australian arid zone: Exploratory time-series analysis of archaeological records. Chungara, 40:245–259, 2008. doi: 10.4067/S0717-73562008000300003.

[120] E. Yáñez, N. A. Lagos, R. Norambuena, C. Silva, J. Letelier, K.-P. Muck, G. S. Martin, S. Benítez, B. R. Broitman, H. Contreras, C. Duarte, S. Gelcich, F. A. Labra, M. A. Lardies, P. H. Manríquez, P. A. Quijón, L. Ramajo, E. González, R. Molina, A. Gómez, L. Soto, A. Montecino, M. Ángela Barbieri, F. Plaza, F. Sánchez, A. Aranis, C. Bernal, and G. Böhm. Impacts of Climate Change on Marine Fisheries and Aquaculture in Chile, volume I, pages 239–332. Wiley-Blackwell, 2017. ISBN 9781119154051. doi: 10.1002/9781119154051.ch10. URL http://doi.wiley.com/10.1002/9781119154051.ch10.

[121] A. F. J. Zangrando. Shell Middens and Coastal Archaeology in Southern South America, pages 9640–9654. Springer International Publishing, 2018. ISBN 978-3-319-51726-1. doi: 10.1007/978-3-319-51726-13024-1”.

