## supplemental material for "Human exploitation of shellfish in the Atacama desert coast and environmental variability: a trans-Holocene perspective"

### Appendix 1: Environmental change and species composition of coastal shellfish assemblages: A past and present comparative perspective on human choices

Bernardo R. Broitman[1,2], Laura Olguín[3], Javiera Guardia[4], Mauricio H. Oróstica[5], Adrien Chevallier[6], Luna Vásquez[7], and Carola Flores[1,8]

[1]Dpto. de Ciencias, Fac. de Artes Liberales, Universidad Adolfo Ibáñez, Viña del Mar, Chile
[2]Instituto Milenio en Socio-ecología Costera (SECOS)
[3]Programa de Doctorado en Antropología UCN-UTA, Universidad Católica del Norte, San Pedro de Atacama, Chile
[4]Universidad SEK, Santiago Chile
[5]Centro de Investigación de Estudios Avanzados del Maule (CIEAM), Universidad Católica del Maule, Talca, Chile
[6]UMR MARBEC, Université de Montpellier, [IFREMER-IRD-CNRS], Montpellier, France
[7]Departamento de Antropología, Facultad de Ciencias Sociales, Universidad de Chile, Santiago, Chile.
[8]Escuela de Arqueología, Universidad Austral de Chile, Puerto Montt, Chile

November 17, 2025

| **Excavation** | I | II | III | IV | V | VI |
|---|---|---|---|---|---|---|
| Zapatero | - | - | 0.49 | - | 0.2 | 0.08 |
| Paposo Norte 09 | 0.1 | - | | 0.09 | - | 0.01 |
| Agua Dulce | - | - | 0.25 | - | - | 0.08 |
| Alero 224A | 0.11 | - | - | 0.01 | - | 0.05 |
| Morro Colorado | - | 0.15 | 0.21 | - | - | |
| Plaza de Indios Norte | - | - | - | - | - | 0.09 |

Table 1: Excavated volumes ($m^2$) from column samples uses for correction of MNI values for each excavation site at every chronocultural period. Site locations are provided in Figure 1 and Table 1 in the main text. A single dash (-) indicates that the chronocultural period was not present in the column sample collected at the site.

| | **SPECIES** | I | II | III | IV | V | VI | Modern |
|---|---|---|---|---|---|---|---|---|
| 1 | Acanthina monodon | 0 | 0 | 2.04 | 0 | 0 | 0 | 0 |
| 2 | Acanthocyclus gayi | 0 | 0 | 0 | 0 | 0 | 0 | 0.08 |
| 3 | Aesopus aliciae | 0 | 0 | 4 | 200 | 0 | 25 | 0 |
| 4 | Argopecten purpuratus | 0 | 20 | 84.19 | 0 | 0 | 23.61 | 0 |
| 5 | Austrolittorina araucana | 0 | 0 | 0 | 0 | 0 | 0 | 0.16 |
| 6 | Austromegabalanus psittacus | 9.09 | 0 | 37.85 | 100 | 0 | 77.5 | 0 |
| 7 | Balanus laevis | 0 | 6.67 | 23.05 | 0 | 0 | 72.5 | 0 |
| 8 | Balanus sp. | 10 | 0 | 4 | 0 | 10 | 99.17 | 0 |
| 9 | Bittium peruvianum | 0 | 0 | 0 | 0 | 0 | 50 | 0 |
| 10 | Brachidontes granulatus | 0 | 0 | 0 | 0 | 0 | 36.11 | 0 |
| 11 | Calyptraeidae | 0 | 0 | 0 | 0 | 0 | 98.61 | 0 |
| 12 | Chiton barnesii | 0 | 0 | 0 | 0 | 0 | 0 | 11.32 |
| 13 | Chiton cumingsii | 0 | 0 | 0 | 0 | 0 | 0 | 2.83 |
| 14 | Chiton granosus | 10 | 26.67 | 304.71 | 900 | 65 | 902.22 | 24.42 |
| 15 | Chiton magnificus | 48.18 | 26.67 | 282.01 | 733.33 | 55 | 624.72 | 0.84 |
| 16 | Chiton sp. | 117.27 | 20 | 291.48 | 222.22 | 15 | 461.67 | 0.19 |
| 17 | Choromytilus chorus | 0 | 26.67 | 94.83 | 11.11 | 0 | 47.22 | 0 |
| 18 | Concholepas concholepas | 160 | 173.33 | 1953.06 | 1855.56 | 65 | 4258.61 | 7.92 |
| 19 | Crassilabrum crassilabrum | 0 | 0 | 14.29 | 0 | 0 | 11.11 | 0 |
| 20 | Crepipatella dilatata | 0 | 0 | 4.76 | 0 | 0 | 0 | 0 |
| 21 | Cyclostremiscus sp. | 0 | 0 | 0 | 0 | 85 | 0 | 0 |
| 22 | Decapoda | 10 | 20 | 58.26 | 511.11 | 15 | 384.17 | 0 |
| 23 | Diloma nigerrimum | 60 | 13.33 | 820.08 | 2244.44 | 510 | 1465.28 | 0 |
| 24 | Echinolittorina peruviana | 20 | 20 | 68.63 | 466.67 | 85 | 1353.61 | 0 |
| 25 | Enoplochiton echinatus | 151.82 | 260 | 1473.25 | 4588.89 | 140 | 5198.61 | 10.79 |
| 26 | Enoplochiton niger | 588.18 | 20 | 509.06 | 1777.78 | 195 | 1304.72 | 66.07 |
| 27 | Felicioliva peruviana | 0 | 20 | 41.93 | 0 | 0 | 0 | 0 |
| 28 | Fissurella bridgesii | 0 | 0 | 23.13 | 0 | 0 | 0 | 0 |
| 29 | Fissurella costata | 0 | 53.33 | 558.91 | 33.33 | 25 | 215.83 | 2.69 |
| 30 | Fissurella crassa | 223.64 | 20 | 1177.09 | 10233.33 | 160 | 4355.56 | 15.11 |
| 31 | Fissurella cumingi | 0 | 0 | 0 | 0 | 0 | 0 | 0.57 |
| 32 | Fissurella latimarginata | 0 | 13.33 | 23.81 | 0 | 0 | 20 | 0.61 |
| 33 | Fissurella limbata | 164.55 | 26.67 | 1869.66 | 777.78 | 140 | 1709.72 | 7.65 |
| 34 | Fissurella maxima | 210 | 220 | 781.61 | 2877.78 | 5 | 956.11 | 5.03 |
| 35 | Fissurella picta | 0 | 0 | 152 | 0 | 0 | 62.5 | 0.83 |
| 36 | Fissurella pulchra | 0 | 0 | 8 | 0 | 0 | 0 | 0 |
| 37 | Fissurella sp. | 202.73 | 106.67 | 687.89 | 1411.11 | 490 | 14691.11 | 0.08 |
| 38 | Heliaster helianthus | 0 | 0 | 0 | 0 | 0 | 0 | 18.67 |
| 39 | Incatella cingulata | 0 | 13.33 | 124.33 | 0 | 10 | 641.67 | 0 |
| 40 | Leptograpsus variegatus | 0 | 0 | 0 | 0 | 0 | 0 | 0.11 |
| 41 | Leukoma thaca | 9.09 | 20 | 139.13 | 0 | 0 | 25 | 0 |
| 42 | Lottia orbigny | 0 | 0 | 0 | 0 | 0 | 0 | 0.2 |
| 43 | Loxechinus albus | 37.27 | 33.33 | 443.13 | 1111.11 | 265 | 3416.67 | 81.92 |
| 44 | Marinula pepita | 20 | 0 | 36.08 | 0 | 0 | 145 | 0 |
| 45 | Mesodesma donacium | 0 | 0 | 2.04 | 0 | 0 | 0 | 0 |
| 46 | Meyenaster gelatinosus | 0 | 0 | 0 | 0 | 0 | 0 | 0.12 |
| 47 | Mitrella sp. | 0 | 0 | 40.76 | 0 | 0 | 100 | 0 |
| 48 | Nassarius gayii | 0 | 33.33 | 55.62 | 0 | 0 | 262.5 | 0 |
| 49 | Perumytilus purpuratus | 0 | 33.33 | 128.57 | 11.11 | 0 | 252.5 | 0 |
| 50 | Petrolisthes violaceus | 0 | 0 | 0 | 0 | 0 | 0 | 0.05 |
| 51 | Priene scabrum | 0 | 6.67 | 4.76 | 0 | 0 | 0 | 0 |
| 52 | Prisogaster niger | 10 | 93.33 | 503.05 | 11.11 | 5 | 300 | 0 |
| 53 | Rissoina inca | 0 | 0 | 24.76 | 0 | 0 | 150 | 0 |
| 54 | Scurria scurra | 0 | 0 | 48.14 | 11.11 | 15 | 12.5 | 0 |
| 55 | Scurria sp. | 495.45 | 46.67 | 649.01 | 3577.78 | 625 | 3764.17 | 12.65 |
| 56 | Scurria viridula | 0 | 0 | 95.16 | 100 | 0 | 195 | 35.77 |
| 57 | Siphonaria lessoni | 0 | 0 | 0 | 0 | 0 | 0 | 1.86 |
| 58 | Stichaster striatus | 0 | 0 | 0 | 0 | 0 | 0 | 7.3 |
| 59 | Taliepus dentatus | 0 | 0 | 0 | 0 | 0 | 0 | 0.12 |
| 60 | Tegula atra | 100 | 2886.67 | 23166.18 | 322.22 | 105 | 10244.44 | 125.03 |
| 61 | Tegula sp. | 196.36 | 0 | 7512 | 8300 | 3565 | 7977.22 | 0 |
| 62 | Tegula tridentata | 0 | 0 | 78.29 | 100 | 0 | 32.5 | 0 |
| 63 | Tetrapygus niger | 0 | 0 | 0 | 0 | 0 | 0 | 52.22 |
| 64 | Thais sp. | 0 | 0 | 14.29 | 0 | 0 | 0 | 0 |
| 65 | Tonicia sp. | 9.09 | 26.67 | 195.56 | 200 | 5 | 243.61 | 3.51 |
| 66 | Trochidae | 27.27 | 0 | 0 | 1400 | 0 | 1220 | 0 |
| 67 | Trochita trochiformis | 9.09 | 0 | 28.76 | 0 | 0 | 25 | 0 |
| 68 | Venerida | 0 | 0 | 16 | 0 | 0 | 37.5 | 0 |
| 69 | Xanthochorus cassidiformis | 0 | 0 | 14.29 | 0 | 0 | 12.5 | 0 |

Table 2: Abundance for all species in the archaeological record (MNI) for each Archaic period (I to VI) and ecological field surveys (individuals per $m^2$). For simplicity, species are listed in alphabetical order.

| **SST** | CDIC | WAIC | EAIC | EBIC | AIC at p.m. | BIC at p.m. | MLL at p.m |
|---|---|---|---|---|---|---|---|
| max | 1302.48 | 743.57 | 842.73 | 1002.58 | 923.61 | 1046.23 | -405.81 |
| min | 1110.08 | 762.06 | 855.32 | 1015.17 | 905.59 | 1028.21 | -396.80 |
| mean | 1147.2 | **706.31** | **805.79** | **965.63** | **452.65** | **575.27** | **-170.33** |
| max, min | 1024.81 | 750.67 | 870.58 | 1054.51 | 935.87 | 1082.58 | -400.94 |
| mean, max | 1293.06 | 733.37 | 854.86 | 1038.79 | 864.33 | 1011.04 | -365.16 |
| mean, min | 1071.15 | 762.53 | 886.20 | 1070.13 | 1041.94 | 1188.65 | -453.97 |
| mean, max, min | **826.97** | 707.77 | 851.51 | 1059.53 | 821.81 | 992.60 | -332.90 |

Table 3: Information criteria obtained after we fitted the latent variable models using all possible combinations of SST statistics, namely the mean and the low and high temperature ranges using the 5, 50, and $95^{th}$ percentiles of all the SST data available for each period for de Holocene shellfish data (min, mean and max, respectively). Abbreviations: CDIC: Conditional Deviance Information Criterion; WAIC: Widely Applicable Information Criterion; post media; EAIC: Expected A Information Criterion; EBIC: Expected Bayesian Information Criterion MLL: Marginal log-likelihood; p.m.: post median. The lowest values are indicated in boldface.

| **SST** | CDIC | WAIC | EAIC | EBIC | AIC at p.m. | BIC at p.m. | MLL at p.m |
|---|---|---|---|---|---|---|---|
| max | 1430.788 | 1032.933 | 1138.859 | 1364.494 | 1100.234 | 1198.100 | -514.117 |
| min | 1398.650 | **1021.418** | **1129.17** | **1354.805** | **1086.59** | **1184.456** | **-507.295** |
| mean | **1325.253** | 1031.873 | 1138.71 | 1364.345 | 1096.551 | 1194.417 | -512.276 |
| maxmin | 1428.462 | 1033.929 | 1150.877 | 1395.542 | 1108.5419 | 1225.436 | -511.27 |
| meanmax | 1450.675 | 1037.321 | 1154.662 | 1399.327 | 1147.178 | 1264.074 | -530.589 |
| meanmin | 1381.437 | 1031.08 | 1151.454 | 1396.119 | 1108.994 | 1225.888 | -511.496 |
| meanmaxmin | 1397.591 | 1032.977 | 1163.919 | 1427.614 | 1483.56 | 1619.488 | -691.782 |

Table 4: Information criteria obtained after we fitted the latent variable models using all possible combinations of SST statistics, namely the mean and the low and high temperature ranges using the 5, 50, and $95^{th}$ percentiles of all the SST data available for each year for the modern fisheries data (min, mean and max, respectively). Abbreviations: CDIC: Conditional Deviance Information Criterion; WAIC: Widely Applicable Information Criterion; post media; EAIC: Expected A Information Criterion; EBIC: Expected Bayesian Information Criterion MLL: Marginal log-likelihood; p.m.: post median. The lowest values are indicated in boldface.
